# Common binding by redundant group B Sox proteins is evolutionarily conserved in *Drosophila*

**DOI:** 10.1101/012872

**Authors:** Sarah H. Carl, Steven Russell

## Abstract

**Background:** Group B Sox proteins are a highly conserved group of transcription factors that act extensively to coordinate nervous system development in higher metazoans while showing both co-expression and functional redundancy across a broad group of taxa. In *Drosophila melanogaster*, the two group B Sox proteins Dichaete and SoxNeuro show widespread common binding across the genome. While some instances of functional compensation have been observed in *Drosophila*, the function of common binding and the extent of its evolutionary conservation is not known.

**Results:** We used DamID-seq to examine the genome-wide binding patterns of Dichaete and SoxNeuro in four species of *Drosophila*. Through a quantitative comparison of Dichaete binding, we evaluated the rate of binding site turnover across the genome as well as at specific functional sites. We also examined the presence of Sox motifs within binding intervals and the correlation between sequence conservation and binding conservation. To determine whether common binding between Dichaete and SoxNeuro is conserved, we performed a detailed analysis of the binding patterns of both factors in two species.

**Conclusion:** We find that, while the regulatory networks driven by Dichaete and SoxNeuro are largely conserved across the drosophilids studied, binding site turnover is widespread and correlated with phylogenetic distance. Nonetheless, binding is preferentially conserved at known cis-regulatory modules and core, independently verified binding sites. We observed the strongest binding conservation at sites that are commonly bound by Dichaete and SoxNeuro, suggesting that these sites are functionally important. Our analysis provides insights into the evolution of group B Sox function, highlighting the specific conservation of shared binding sites and suggesting alternative sources of neofunctionalisation between paralogous family members.

## Background

The importance of regulatory DNA in development, evolution and disease has become widely recognized in recent years as the number of functional genomic studies mapping regulatory elements in the genomes of humans and model organisms has grown [1–5]. While significant progress has been made on questions such as the *in vivo* binding specificity of transcription factors (TFs) and the combinatorial patterns of TF binding that drive specific gene expression, in many cases the large number of binding events observed *in vivo* suggests that TF function is context-dependent and influenced by factors in addition to their intrinsic sequence specificity [6–12]. Furthermore, the most common methods of assaying *in vivo* genome-wide binding are noisy and may suffer problems of bias and false positives [13–17]. One way to cut through this noise and assess the functionality of TF binding is to leverage the effect of natural selection, which should tend to preserve functionally important binding events during evolution [6, 18–20]. Here we have used a comparative evolutionary approach in four species of *Drosophila* to study the binding and functions of two group B Sox proteins, a family of TFs that is both deeply conserved throughout animal evolution and displays evidence of functional redundancy.

*Sox* (SRY-related high-mobility group box) genes encode a highly conserved family of TFs that serve as broad developmental regulators in metazoa. They are thought to have evolved in conjunction with the origin of multicellular animal life, as they are present in all animal genomes in which they have been searched for, including basal members such as sponges and placozoa [21–25]. All Sox proteins contain a single HMG (high-mobility group) DNA binding domain, and they are subdivided into ten groups, A through J, based on HMG sequence and full-length protein structure [24, 26–28]. In vertebrates, members of each group are often expressed in overlapping patterns in particular tissues during development and play important roles in directing the differentiation of cells in those tissues. For example, group B genes are expressed in the developing central nervous system (CNS) and eye [29–32], group C genes are expressed in the kidney and pancreas [33–35], and group F genes are expressed in the developing vascular and lymphatic system [36, 37]. Interestingly, not only are group members co-expressed, in many cases they show a pattern of phenotypic redundancy where severe developmental defects are only observed when multiple members of the same group are removed [27, 37–43]. While mammalian genomes contain multiple paralogues for most groups, invertebrates typically have fewer Sox genes. Sequenced insect genomes, including that of the fruit fly *Drosophila melanogaster*, typically contain one gene in each of groups C, D, E and F, and four genes in group B, although occasional extra genes are observed in certain lineages [24, 26].

Group B *Sox* genes are of particular evolutionary interest because, in addition to being the most closely related *Sox* genes to *Sry*, they are some of the best characterized members of the *Sox* family and appear to have highly conserved functions throughout evolution [44, 45]. In mammals, group B Sox genes are involved in stem cell pluripotency and self-renewal, ectoderm formation, neural induction, CNS development, placode formation and gametogenesis [27]. Vertebrate group B Sox genes are divided into two subgroups: group B1, which includes *Soxl, Sox2* and *Sox3* [44], and group B2, which includes *Sox14* and *Sox21* [46]. Group B1 and B2 genes play opposing roles in the developing vertebrate CNS, with group B1 proteins primarily acting as transcriptional activators to convey early neuroectodermal competence and maintain neural precursors while group B2 genes primarily act as transcriptional repressors to promote neuronal differentiation [43, 47–49]. Although functional data suggests that the B1-B2 division may not be an appropriate classification in insects [45, 50, 51], a role for group B Sox genes in neural development appears to be conserved, making *Drosophila* an attractive system in which to study group B *Sox* function and evolution more closely [31, 32, 43]. There are four group B *Sox* genes in the *Drosophila melanogaster* genome: *SoxNeuro (SoxN), Dichaete, Sox21a* and *Sox21b*. Of these, the best studied to date are *Dichaete* and *SoxN*. While the exact orthology relationships between vertebrate and insect group B *Sox* genes are still debated, similarities in the expression patterns and functions of *Soxl*, *Sox2* and *Sox3* in vertebrates and *SoxN* and *Dichaete* in insects suggest a deeply conserved yet complex relationship between these two sets of Sox genes [31, 32, 45, 50, 52–60].

As with *Soxl, Sox2* and *Sox3* in vertebrates, both *Dichaete* and *SoxN* are expressed in overlapping patterns in the developing *Drosophila* CNS and are necessary for its normal development; they also show evidence of functional redundancy [29, 55, 61–64]. *Dichaete* mutant embryos show axonal and midline defects, which can be rescued by expressing Dichaete or mammalian Sox2 in the midline [63]. *SoxN* mutant embryos also show axonal defects and loss of lateral neurons that can be rescued with mammalian Soxl [65]. *SoxN/Dichaete* double mutants show much more severe CNS defects than either single mutant, in particular an increased loss of neuroblasts in the medial and intermediate columns of the neuroectoderm, where *SoxN* and *Dichaete* expression overlaps most strongly [61, 66]. In addition to compensation at the level of neural phenotypes, *in vivo* binding and expression studies of Dichaete and *SoxN* in *D. melanogaster* have shown that they have highly similar genome-wide binding patterns and share a large number of target genes [51, 67]. Commonly bound targets cover many of the core functions of both Dichaete and SoxN, including over a hundred other TFs active in the CNS, the proneural genes of the achaete-scute complex, *Drop* (*Dr*) and *ventral nervous system defective* (*vnd*), which encode TFs involved in CNS dorso-ventral patterning [68], and the neuroblast temporal identity genes *seven-up* (*svp*), *hunchback* (*hb*), *Kruppel* (*Kr*) and *POU domain protein 2* (*pdm2*) [51, 67, 69, 70].

Not only do Dichaete and SoxN share many targets, they also display a complex pattern of compensatory binding in each other’s absence. Studies measuring SoxN binding in *Dichaete* mutants and *vice versa* identified loci where one TF can compensate for the other’s absence by binding in its place or increasing its own binding. However, at other loci, the loss of one Sox protein appears to result in a loss of binding by the other. These observations suggest that in some circumstances Dichaete and SoxN can compensate for one another, but in other case they are dependent on one another. Furthermore, at certain loci their binding is independent, with the loss of one TF not affecting the binding of the other. It has been suggested that this complex interplay may be the result of partial neofunctionalisation between the two paralogous TFs leading to their unique roles, while maintaining some degree of redundancy in order to provide robustness to the critical process of early neural development [51, 71–73]. However, it is not known whether the overlap in binding patterns and targets observed between Dichaete and SoxN, which is substantially greater than that observed between many other paralogous genes such as the *cis*-paralogous members of the same *Hox* clusters [7, 8, 11, 74, 75], has been specifically maintained during evolution. Given the fact that functional redundancy among *Sox* genes from the same group seems to be a common theme across many different taxa, we were curious to examine the question of group B *Sox* binding from an evolutionary perspective.

Comparative studies of transcription factor binding can facilitate an analysis of the evolutionary dynamics of regulatory DNA as well the use of natural selection as a filter to reduce the biological and technical noise inherent to *in vivo* genome-wide binding assays [6, 15, 76–78]. Previous comparative studies in *Drosophila* have primarily used ChIP-seq to measure binding and have focused on developmental regulators such as the anterioposterior (AP) patterning factors Bicoid (Bcd), Hunchback (Hb), Kruppel (Kr), Giant (Gt), Knirps (Kni) and Caudal (Cad) and the mesodermal regulator Twist (Twi) [76, 77, 79]. These studies measured both quantitative turnover of binding sites during evolution as well as qualitative changes in binding strength, finding broadly similar rates of divergence for different factors. They also used the evolutionary data to discover features of TF binding that were associated with higher conservation, such as clustered binding, combinatorial binding with other TFs, and the presence of specific sequence motifs in or near binding intervals [76, 77]. The degree of conservation of binding events between different *Drosophila* species, which is generally greater than that between equally distant vertebrate species [20, 80–82], makes *Drosophila* a particularly well-suited model system for studying the evolution of regulatory DNA and for making inter-species comparisons of TF binding. With this in mind, we used DamID-seq to study the *in vivo* binding patterns of Dichaete and SoxN in four species of *Drosophila*, *D. melanogaster*, *D. simulans*, *D. yakuba* and *D. pseudoobscura*, with the goal of shedding new light on the functional and evolutionary dynamics of group B Sox binding.

## Results

### Comparative DamID-seq for group B Sox proteins

We have recently used combinations of genome-wide ChIP and DamID data to define high confidence *in vivo* binding profiles and core binding intervals for both Dichaete and SoxN in *Drosophila melanogaster* [51, 67]. In order to better understand aspects of group B Sox functional compensation at a genomic level, we expanded on this work, performing DamID using *D. melanogaster* Sox-Dam fusion proteins and high-throughput sequencing to map Dichaete binding in four *Drosophila* species and SoxN binding in two species (Figure 1). The amino acid sequences of both proteins are highly conserved across the four species; thus our experimental design facilitates an analysis of binding attributable to differences in the genome sequence or nuclear environment between species, rather than to the Sox proteins themselves (Supplementary Figure 1). We used a differential enrichment approach to identify GATC fragments in each genome that were significantly bound by each Sox protein (Dichaete-Dam or SoxN-Dam) in comparison to Dam-only controls and merged neighbouring fragments to generate binding intervals (Table 1A). While these binding intervals do not precisely correspond to ChIP-seq peaks or DamID peaks we previously determined via tiling microarrays, we nevertheless found that, in agreement with previous studies, both Dichaete and SoxN showed widespread binding at thousands of sites across each fly genome [51, 67]. After normalisation and correction for multiple hypothesis testing, we identified between 17,000 – 26,000 binding intervals (p < 0.05) for Dichaete in *D. melanogaster, D. simulans* and *D. yakuba*, and for SoxN in *D. melanogaster* and *D. simulans*. Approximately 3000 Dichaete binding intervals were identified in *D. pseudoobscura*, reflecting the fact that the binding profiles were noisier in this species compared to the other three species, with less reproducible biological replicates.

**Figure 1.**
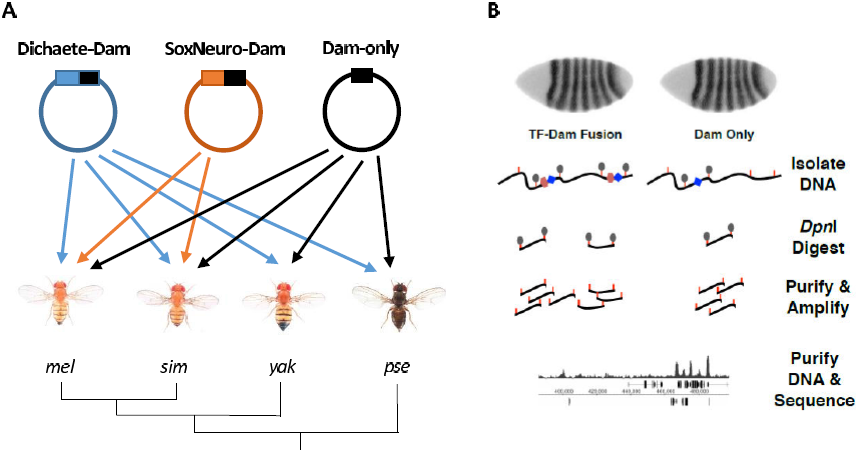
Overview of experimental strategies. (A) Schematic of transgenic lines generated in each species of *Drosophila*. Three plasmids, containing either a Dichaete-Dam fusion protein, a SoxN-Dam fusion protein, or Dam only, were injected. Dichaete-Dam and Dam-only lines were created for each species, while SoxN-Dam lines were only created in *D. melanogaster* and *D. simulans*. The dendrogram represents the evolutionary relationships between the species used. Photographs of flies are from N. Gompel (http://www.ibdml.univ-mrs.fr/equipes/BPNG/Illustrations/melanogaster%20subgroup.html). Abbreviations: *mel, Drosophila melanogaster; sim, Drosophila simulans; yak, Drosophila yakuba; pse, Drosophila pseudoobscura*. (B) Outline of the DamID-seq experimental protocol. A TF-Dam fusion protein or a Dam-only control is expressed in embryos, leading to methylation of GATC sites in the vicinity of binding events. Genomic DNA is extracted and methylated fragments are isolated via digestion with *Dpn*I, which recognises only methylated GATC. These fragments are purified through ligation of adapters, further digestion of non-methylated DNA with *Dpn*II and PCR amplification [95]. DNA from both TF-Dam fusion samples and control Dam-only samples is sequenced and mapped to the genome, and the relative enrichment of reads is compared between conditions to generate binding profiles. Orange lines represent GATC sites, grey ovals represent methylation, blue diamonds represent Dam protein and pink ovals represent TFs. Figure adapted from Carl and Russell (in press).

**Table 1.** Binding intervals found for each DamID dataset and genes annotated to intervals.

| **A** | Binding intervals<br>($p < 0.01$) | Binding intervals<br>($p < 0.05$) |
| --- | --- | --- |
| *D. mel* D-Dam | 17530 | 20848 |
| *D. mel* SoxN-Dam | 17833 | 22952 |
| *D. sim* D-Dam | NA | 17833 |
| *D. sim* SoxN-Dam | NA | 17209 |
| *D. yak* D-Dam | 21988 | 26563 |
| *D. pse* SoxN-Dam | NA | 2951 |

| **B** | Translated binding intervals | Overlaps with *D. mel* binding intervals | % of *D. mel* intervals overlapping | % of non-*D. mel* intervals overlapping |
| --- | --- | --- | --- | --- |
| *D. sim* D-Dam | 16119 | 11647 | 55.9 | 72.3 |
| *D. sim* SoxN-Dam | 15142 | 11891 | 51.8 | 78.5 |
| *D. yak* D-Dam | 20964 | 14573 | 69.9 | 69.5 |
| *D. pse* D-Dam | 2020 | 1301 | 6.24 | 64.4 |

| **C** | Genes annotated to binding intervals | Genes common to Dichaete and SoxN | Direct targets annotated to binding intervals |
| --- | --- | --- | --- |
| *D. mel* D-Dam | 9400 | 8445 | 1173/1373 (85.4%) |
| *D. mel* SoxN-Dam | 9528 | 8445 | 434/544 (80.0%) |
| *D. sim* D-Dam | 9383 | 7524 | 1111/1373 (80.9%) |
| *D. sim* SoxN-Dam | 8948 | 7524 | 412/544 (75.7%) |
| *D. yak* D-Dam | 12192 | NA | 1249/1373 (91.0%) |
| *D. pse* D-Dam | 1888 | NA | 407/1373 (29.6%) |

Translating sequence reads from each non-melanogaster species to the *D. melanogaster* genome coordinates and calling binding intervals on the translated data showed that, while differences between species were observed, many binding intervals were conserved across the species (Figure 2A-2B; Table 1B). Additionally, and in agreement with our previous work, the Dichaete and SoxN binding profiles showed a high degree of concordance in both *D. melanogaster* and *D. simulans* [51]. We used this translated binding data to assess the gene targets and genomic features associated with each dataset. Assigning each binding interval to a putative target gene identified approximately 9000-9500 genes associated with both Dichaete and SoxN binding in *D. melanogaster* and *D. simulans* (Table 1C; Additional File 1). A high proportion of these genes are shared between Dichaete and SoxN in both species. Dichaete binding intervals in *D. yakuba* were associated with over 12,000 genes, while in *D. pseudoobscura* we identified 1888 associated genes. With the exception of the *D. pseudoobscura* data, each set of putative target genes contains a high percentage of previously identified Dichaete or SoxN direct target genes (Table 1C) [51, 67]. In line with previous studies of Dichaete and SoxN binding, all sets of target genes are highly enriched for specific Gene Ontology Biological Process (GO:BP) terms such as generation of neurons (p < 1e-29) and regulation of transcription (p < 1e-9) [51, 67] as well as higher level terms such as general organ and system development (p < 1e-47) and biological regulation (p < 1e-44) (Additional File 2). Notably, although there were many fewer Dichaete bound genes identified in *D. pseudoobscura*, these show strong enrichment for similar GO:BP terms, are strongly upregulated in the brain and larval CNS, and are highly associated with publications describing genes involved in the neural stem cell transcriptional network (p < 1e-25), all of which are hallmarks of known Dichaete functions [50, 64, 67]. We also used the translated binding data to determine the location of binding intervals with respect to transcription start sites. In broad agreement with our previous work [51, 67], we found similar distributions in each species: approximately 65% of binding intervals overlap introns, with the remainder mostly mapping in the proximity of promoters and a minority within intergenic regions.

**Figure 2.**
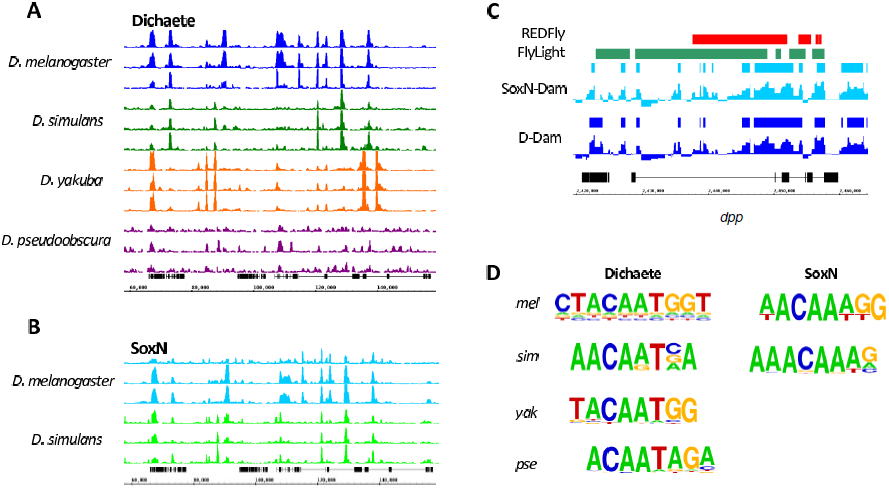
Results of comparative DamID for Dichaete and SoxN. (A) Dichaete-Dam binding profiles plotted on the *D. melanogaster* genome showing translated reads mapping to an orthologous region in four species. Three biological replicates from each species are shown. All read libraries were scaled to a total size of 1 million reads for visualization purposes. The y-axes of all tracks range from 0-50 reads. (B) SoxN-Dam binding profiles plotted on the *D. melanogaster* genome showing translated reads mapping to an orthologous region in two species. Three biological replicates from each species are shown. All read libraries were scaled to a total size of 1 million reads for visualization purposes. The y-axes of all tracks range from 0-50 reads. The same ~120-kb region of chromosome 2L is shown in (A) and (B). (C) Overlaps between *D. melanogaster* Sox DamID binding intervals and known CRMs from REDFly and FlyLight at the *decapentaplegic (dpp)* locus. Binding profiles represent the normalised log2 fold changes between Sox fusion binding and Dam-only control binding in each GATC fragment. (D) *De novo* Sox motifs discovered in DamID binding intervals. Motifs discovered in Dichaete intervals in each species show a preference for T in the fourth position of the core CAAAG motif, while those discovered in SoxN intervals in each species show a preference for A. Abbreviations: *mel, Drosophila melanogaster; sim, Drosophila simulans; yak, Drosophila yakuba; pse, Drosophila pseudoobscura*.

We performed searches for known and *de novo* binding motifs with the binding intervals in each original genome to identify any differences in preferential motif usage between species or TFs. The known motif searches uncovered highly significant matches to vertebrate Sox2, Sox3 and Sox6 motifs. *De novo* searches found a significantly enriched motif matching the consensus Sox motif in each dataset (p <= 1e-20), which correspond well to motifs discovered in our previous Dichaete and SoxN studies [51, 67](Additional File 3). Of particular interest, we found that the Sox motifs identified in the Dichaete binding intervals of each species differed from those found for SoxN (Figure 2D). The primary difference is in the fourth position of the core CAAAG motif, which shows a stronger preference for T in the Dichaete intervals and for A in the SoxN-Dam intervals from each species. Although it is not known whether these differences in Sox motifs found in Dichaete and SoxN binding intervals are responsible for different functions of the two TFs, it is striking that the same patterns appear independently in the genomes of multiple species. In addition, we found a set of motifs associated with a broader array of TFs: in the case of Dichaete these include known interaction partners Ventral veins lacking (Vvl) and Nubbin (Nub) as well as the early segmentation factors Knirps (Kni) and Runt (Run).

Finally, we examined the overlap between group B Sox binding intervals and known *cis*-regulatory modules (CRMs) by comparing our *D. melanogaster* data with annotated CRMs from the REDFly and FlyLight databases as well as a recent STARR-seq assay identifying active enhancers [83–85]. Both Dichaete and SoxN show high overlap with CRMs from all three sources, with a high proportion of CRMs overlapping both Dichaete and SoxN binding intervals (Table 2). The highest overlap was found with the REDFly database (~60% of CRMs); while the FlyLight and STARR-seq data showed slightly lower overlap, in the case of FlyLight we found higher overlap with CNS specific CRMs. Although the intervals mapped with DamID do not correspond directly to enhancer elements, in many cases visual inspection reveals a very good correlation between annotated CRMs and peaks of DamID binding (e.g. *dpp*, Figure 2C). Together with the gene and genomic feature annotations, these observations support the emerging view that Dichaete and SoxN work together to regulate a large set of genes involved in CNS development through binding at known CRMs, with a preference for intronic binding [51, 67], and suggest that the overall roles of these group B Sox proteins have not changed significantly during the evolution of the drosophilids analysed here.

**Table 2.** Overlaps between Sox binding intervals and known CRMs.

| Dataset | CRMs in dataset | CRMs overlapping Dichaete binding | CRMs overlapping SoxN binding | CRMs overlapping Dichaete and SoxN binding |
| --- | --- | --- | --- | --- |
| REDFly | 1864 | 1152 (61.8%) | 1130 (60.6%) | 1108 (59.4%) |
| FlyLight | 7113 | 2999 (42.2%) | 2784 (39.1%) | 2681 (37.8%) |
| FlyLight CNS | 4727 | 2077 (44.0%) | 1935 (41.0%) | 1857 (39.3%) |
| STARR-seq S2 | 2325 | 1092 (47.0%) | 951 (40.9%) | 912 (39.2%) |
| STARR-seq OSC | 3341 | 1144 (34.2%) | 1061 (31.8%) | 973 (29.1%) |

### Turnover of Dichaete binding sites during evolution

In order to examine the patterns of group B Sox binding conservation and turnover during evolution, we focused our analysis on the Dichaete-Dam binding data in *D. melanogaster, D. simulans* and *D. yakuba*. While the *overlap* in target genes between these three species is high, we found widespread turnover in the orthologous locations of binding intervals. Indeed, out of all 26,117 Dichaete binding intervals identified across the three species, the major fraction (47%) are present in only one species, with smaller proportions present at orthologous locations in two (23%) or all three (30%) species (Figure 3A). Clustering all Dichaete-Dam replicates by binding affinity scores reveals that the three biological replicates from each species cluster tightly (Pearson’s correlation coefficients from 0.92-0.99, Figure 3B), but replicates from different species also show very good correlations (PCC = 0.64-0.74). The patterns of similarities and differences between Dichaete binding patterns in each species, as visualized by principal component analysis (PCA) on the normalised binding affinity scores in all bound intervals, are consistent with the expectation of neutral evolution along the *Drosophila* phylogeny.

**Figure 3.**
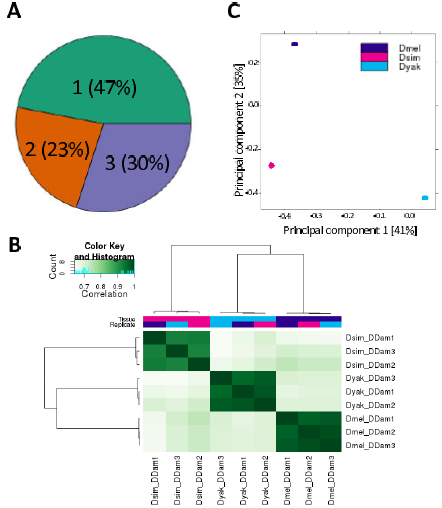
Three-way comparison of Dichaete binding. (A) Pie chart showing the percentage of all Dichaete binding intervals identified that are present in one species (47%), two species (23%) or all three species (30%). (B) Heatmap showing all biological replicates of Dichaete-Dam samples in *D. melanogaster*, *D. simulans* and *D. yakuba* clustered by binding affinity score (log of normalised read counts) within bound intervals. The color key and histogram shows the distribution of correlation coefficients for affinity scores between each pair of samples. Darker green corresponds to a higher correlation between samples, while lighter green corresponds to a lower correlation. Biological replicates from each species cluster strongly together, though good correlations are seen even between species. (C) Plot of principal component analysis (PCA) of Dichaete-Dam binding affinity scores in bound intervals in *D. melanogaster, D. simulans* and *D. yakuba*. The first principal component (x-axis) separates *D. yakuba* from the other two species, while the second principal component (y-axis) separates *D. melanogaster* from *D. simulans* and *D. yakuba*.

We employed DiffBind [86] to perform a differential enrichment analysis, comparing Dichaete-Dam binding profiles between two species for each binding interval, identifying binding intervals that showed significant quantitative differences between each pair of species [86]. For comparisons between *D. melanogaster* and *D. simulans* or *D. yakuba*, we found approximately equal numbers of preferentially bound intervals in each species (Figure 4A-B). In agreement with the PCA plot, 8880 differentially bound intervals were identified between *D. melanogaster* and *D. yakuba* at FDR1, while only 5044 were identified between *D. melanogaster* and *D. simulans*, indicating a greater amount of quantitative binding divergence between *D. melanogaster* and *D. yakuba*. Again, this finding shows that divergence in binding follows the known *Drosophila* phylogeny and suggests a molecular clock mechanism for Dichaete binding site turnover, as has been proposed for other TFs in *Drosophila* [77]. Interestingly, the proportion of differentially bound intervals that are present in both species but show quantitative changes in binding strength, as opposed to those that are qualitatively absent in one species, also increases with phylogenetic distance, from 58.0% between *D. melanogaster* and *D. simulans* to 63.7% between *D. melanogaster* and *D. yakuba*. This also represents an increase in the percentage of the total *D. melanogaster* Dichaete binding intervals that quantitatively change in another species, from 9.6% in *D. simulans* to 20.4% in *D. yakuba*.

**Figure 4.**
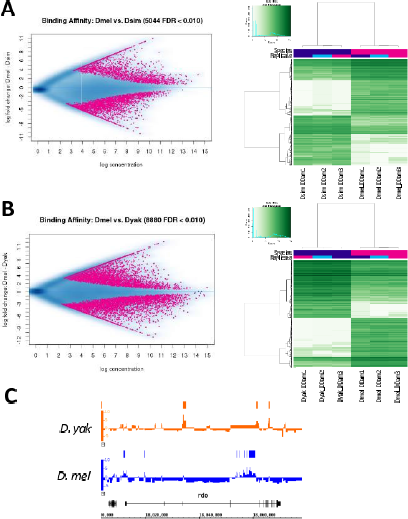
Pairwise analysis of Dichaete binding site turnover. (A) Quantitative differences in Dichaete binding between *D. melanogaster* and *D. simulans*. MA plot shows binding intervals that are preferentially bound in *D. melanogaster* (pink, fold change > O) or *D. simulans* (pink, fold change < 0) at FDR1. Heatmap shows differentially bound Dichaete-Dam intervals between *D. melanogaster* and *D. simulans* clustered by binding affinity scores. The color key and histogram shows the distribution of binding affinity scores (log of normalised read counts) in all bound intervals in each sample. Darker green corresponds to higher affinity scores or stronger binding, while lighter green corresponds to lower affinity scores or weaker binding. Roughly equal numbers of intervals are preferentially bound in each species. (B) Quantitative differences in Dichaete binding between *D. melanogaster* and *D. yakuba*. MA plot shows binding intervals that are preferentially bound in *D. melanogaster* (pink, fold change > 0) or *D. yakuba* (pink, fold change < O) at FDR1. Heatmap shows differentially bound Dichaete-Dam intervals between *D. melanogaster* and *D. yakuba* clustered by binding affinity scores. The color key and histogram are as in (A). Again, rough equal numbers of intervals are preferentially bound in each species, although more differentially bound intervals are identified overall. (C) Example of Dichaete binding site turnover between *D. melanogaster* (blue) and *D. yakuba* (orange) at the *reduced ocelli* (*rdo*) locus. Binding profiles represent the normalised log2 fold changes between Dichaete-Dam binding and Dam-only control binding in each GATC fragment. Bars represent bound intervals identified at FDR5. Bound intervals that are positionally conserved are not shown. Strong binding is observed in the third, fourth and eleventh introns in *D. yakuba*; these binding events are lost in *D. melanogaster*, but several binding intervals are gained in the first and fourth introns.

Under balancing selection it is proposed that, as the frequency of conserved binding events at orthologous positions decreases between more distantly related species, new binding events at the same gene loci should evolve to maintain the same level of gene expression. This is often referred to as binding site turnover or compensatory evolution [19, 76, 77, 83]. In order to detect such events, we considered the set of Dichaete binding intervals in one species that do not overlap with any interval in a pairwise comparison with each other species, as well as the genes to which they are annotated. Binding intervals annotated to the same gene but which do not overlap in orthologous position were considered instances of compensatory evolution. We found instances of binding site turnover between D. melanogaster and D. simulans at 2457 genes, and between D. melanogaster and D. yakuba at 2806 genes; an example of turnover between D. melanogaster and D. yakuba at the rdo locus is shown in Figure 4C. We were curious as to whether these compensatory binding sites are located within CRMs that are active in both species or whether they arise in conjunction with the appearance of new functional CRMs. To address this, we utilised data from STARR-seq assays performed in *D. yakuba* and *D. melanogaster* where it is estimated that 19% of active *D. melanogaster* CRMs show compensatory conservation relative to *D. yakuba* CRMs [83]. In order to take a broad view of Sox binding at CRMs, we re-analysed the low-stringency STARR-seq data and identified 21105 *D. melanogaster* CRMs active in S2 cells that show compensatory conservation relative to *D. yakuba* and 22444 that show compensatory conservation relative to *D. melanogaster*. In ovarian somatic cells (OSCs), we found 12843 *D. melanogaster* CRMs and 20207 *D. yakuba* CRMs that show compensatory conservation. In *D. melanogaster*, only 53 S2 cell and 105 OSC compensatory CRMs contain Dichaete binding intervals that are also compensatory relative to *D. yakuba*, while in *D. yakuba*, only 90 S2 and 157 OSC compensatory CRMs contain Dichaete binding intervals that are also compensatory relative to *D. melanogaster*. This observation strongly suggests that the majority of Dichaete binding site turnover events happen in active CRMs that are functionally and positionally conserved in both species.

### Binding conservation and regulatory function

Focusing on the Dichaete binding intervals that are positionally conserved between species, we assessed whether this type of conservation is enriched at certain classes of functional sites, such as known CRMs or Dichaete target genes. Such an enrichment has previously been found for other TFs in Drosophila, including the AP factors Bcd, Hb, Kr, Gt, Kni and Cad [76] and the mesoderm regulator Twi [77]. We first examined Dichaete binding intervals associated with REDFly and FlyLight CRMs [84, 85] and found that 64.4% of Dichaete binding intervals associated with a REDFly CRM are conserved in all three species, compared to only 44.8% of binding intervals that are not at REDFly CRMs (Table 3). Conversely, only 9.6% of Dichaete intervals at REDFly CRMs are unique to *D. melanogaster*, while 27.6% of those that are outside REDFly CRMs are unique (Supplementary Figure 3). Similar results were found for binding intervals within FlyLight enhancers. Overall, being located at a known CRM has a highly significant effect on Dichaete binding interval conservation (REDFly: χ^2^ = 161.9, d.f. - 3, p = 7.06e-35; FlyLight: χ^2^ = 177.3, d.f. = 3, p-value = 3.38e-38). We also analysed this effect on two-way conservation of SoxN binding intervals between *D. melanogaster* and *D. simulans*, again finding that CRM-associated binding is significantly more likely to be conserved between species (REDFly: χ^2^ = 323.6, d.f. = 1, p = 2.40e-72; FlyLight: χ^2^ = 46.95, d.f. = 1, p = 7.27e-12). The strong increase in rates of conservation observed for group B Sox binding intervals within CRMs suggests that these sites are likely to be under balancing selection to maintain their effect on gene regulation and is consistent with the important regulatory functions of Dichaete and SoxN [51, 67].

**Table 3.** Conservation of Sox binding intervals at functional sites.

| Factor | Condition | Percent of binding intervals conserved | Percent of binding intervals unique to *D. mel* |
| --- | --- | --- | --- |
| Dichaete | REDFly | 64.4% | 9.6% |
|  | Not REDFly | 44.8% | 27.6% |
| SoxN | REDFly | 79.0% | 21.0% |
|  | Not REDFly | 46.5% | 53.5% |
| Dichaete | FlyLight | 54.7% | 16.7% |
|  | Not FlyLight | 44.2% | 28.6% |
| SoxN | FlyLight | 65.3% | 34.7% |
|  | Not FlyLight | 45.1% | 54.9% |
| Dichaete | Core interval | 70.4% | 7.7% |
|  | Not core interval | 38.5% | 32.4% |
| SoxN | Core interval | 75.7% | 24.3% |
|  | Not core interval | 44.7% | 55.3% |
| Dichaete | Direct target | 53.7% | 20.4% |
|  | Not direct target | 43.1% | 29.0% |
| SoxN | Direct target | 53.3% | 47.6% |
|  | Not direct target | 47.3% | 52.7% |

Our previous experiments identified *D. melanogaster* Dichaete and SoxN direct target genes and high-confidence core binding intervals supported by both ChIP and DamID data [51, 67]. Given that Dichaete and SoxN binding intervals in known CRMs are preferentially conserved, indicating a link between functional binding and conservation, we asked whether they are also highly conserved at core intervals and direct target genes. The *D. melanogaster* FDR1 Dichaete-Dam intervals identified in this study show a greater overlap with core Dichaete intervals than the FDR1 SoxN-Dam intervals do with SoxN core intervals. However, for both TFs, DamID binding intervals that overlap a core interval are significantly more likely to be conserved than those that do not (Dichaete: χ^2^ = 1408.6, d.f. = 3, p = 4.10e-305; SoxN: χ^2^ = 733.1, d.f. = 1, p-value = 1.90e-161) (Supplementary Figure 4). For both TFs, this effect is even stronger than that of being located within a known CRM. Although core binding intervals do not necessarily represent direct targets, as measured by gene expression assays, these data show that they are nonetheless subject to evolutionary constraint and are likely to be functionally important. Interestingly, we found that for both Dichaete and SoxN, association with a direct target gene has less of an effect on binding interval conservation than being located at a core interval (Dichaete: χ^2^ = 57.3, d.f. = 3, p = 2.3e-12; SoxN: χ^2^ = 17.2, d.f. = 1, p = 3.4e-5) (Supplementary Figure 4). This is not as surprising as it may first seem since direct targets are identified by expression changes in mutant embryos, and it is clear that Dichaete and SoxN can functionally compensate for each other’s loss, masking significant expression changes at some targets. In addition, in many cases, multiple binding intervals are annotated to the same target gene, and these are unlikely to be equally functional. For example, some may represent shadow enhancers and may therefore be less constrained by natural selection [87, 88]. The presence of these functionally less constrained binding intervals in our datasets is likely to reduce the overall rate of conservation observed for intervals annotated to direct target genes compared to those at core intervals, which are robustly detected by independent binding assays.

### Sequence conservation of Sox motifs and binding intervals

We used the reference genome sequence for each species to assess the contribution to binding conservation of sequence conservation within binding intervals and at TF-specific binding motifs. In order to examine the patterns of motif conservation in Dichaete binding intervals, we first identified all matches to the best *de novo Sox* motif discovered in each set of intervals [89]. We did the same with control binding intervals that had been randomly shuffled to different locations in each genome [90]. In all cases, significantly more Sox motifs were found in Dichaete binding intervals than in control intervals (p < 4.03e-15, Wilcoxon rank sum test with continuity correction). Focusing on Dichaete binding intervals, we compared intervals that are bound in all four species, including *D. pseudoobscura*, to those that are unique to *D. melanogaster*. The highly conserved intervals contain significantly more Sox motifs on average (mean = 4.53) than the binding intervals unique to *D. melanogaster* (mean = 1.29, p = 3.03e-193, Wilcoxon rank sum test with continuity correction), showing that the presence and number of Sox motifs is positively correlated with Dichaete binding conservation (Figure 5A).

**Figure 5.**
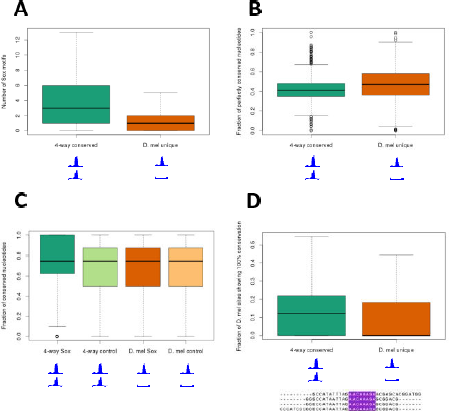
Density and conservation of Sox motifs are higher in Dichaete intervals with binding conservation. (A) On average, Dichaete binding intervals that are conserved between all four species (“4-way conserved”) have more Sox motifs than intervals that are unique to *D. melanogaster* (“D. mel unique”) (p = 3.03e-193). (B) Dichaete binding intervals that are conserved between all four species do not show an increased rate of total nucleotide conservation on average than intervals that are unique to *D. melanogaster*. (C) Sox motifs in Dichaete intervals that are bound in all four species (“4-way Sox”) have a greater percentage of perfectly conserved nucleotides across all species than either Sox motifs in intervals that are unique to *D. melanogaster* (“D. mel Sox”, p = 9.56e-36), randomly shuffled control motifs in intervals that are bound in all four species (“4-way control”, p = 1.62e-43) or randomly shuffled control motifs in intervals that are unique to *D. melanogaster* (“D. mel control”, p = 1.67e-9). (D) On average, Dichaete intervals that are bound in all four species have more Sox motifs that are both positionally conserved and show 100% nucleotide conservation across all species than intervals that are only bound in *D. melanogaster* (p = 6.04e-28). The multiple alignment illustrates a positionally conserved Sox motif with 100% nucleotide conservation (highlighted in purple).

Previous comparative ChIP-seq studies of TF binding in *Drosophila* have found that overall nucleotide conservation is not significantly elevated in binding intervals that are conserved between species [76, 77]. To determine whether this is also the case for our DamID data, we used PRANK, a phylogeny-aware aligner [91, 92], to create multiple alignments of high-confidence orthologous sequences from each species with Dichaete binding intervals showing four-way binding conservation and those showing unique *D. melanogaster* binding. These sequences should contain the regions of regulatory DNA to which Dichaete binds; however, they are also likely to contain flanking regions that are not functionally relevant. Not surprisingly, we did not detect a higher rate of nucleotide conservation in intervals showing four-way binding conservation compared to the unique *D. melanogaster* intervals; in fact, the uniquely-bound intervals show slightly, but significantly, greater sequence conservation across their entire lengths (Wilcoxon rank sum test, p = 2.34e-20) (Figure 5B). We scanned the multiple alignments for matches to the *de novo* Sox motifs we identified and calculated the rates of nucleotide conservation specifically within these motifs [93, 94]. In contrast to the full binding interval sequences, Sox motifs within intervals showing four-way conservation show significantly higher rates of nucleotide conservation than those in intervals that are only bound in *D. melanogaster* (Wilcoxon rank sum test, p = 9.56e-36). As a further control, we randomly shuffled the Sox motifs to produce a set of motifs with the same GC content and length and searched for matches to each of them in the multiply aligned binding intervals. While the average rates of nucleotide conservation in matches to shuffled control motifs are slightly higher in intervals that display four-way binding conservation than in unique *D. melanogaster* intervals, Sox motifs in intervals with four-way binding conservation are significantly more highly conserved than control motifs in either set of intervals (Wilcoxon rank sum test, p = 1.63e-42 and p = 1.67e-9) (Figure 5C).

We also searched for Sox motifs that were independently identified at the exact orthologous location in all four species (positionally conserved). We found that approximately 20% of Sox motifs in binding intervals showing four-way binding conservation are positionally conserved, as opposed to 16% of Sox motifs in intervals that are only bound in *D. melanogaster*, a significant difference (Wilcoxon rank sum test, p = 2.55e-24). A similar pattern holds for the subset of positionally conserved motifs that show complete sequence conservation between all four species (Figure 5D). These perfectly conserved motifs make up approximately 15% of Sox motifs in intervals that show four-way binding conservation but only 12% of Sox motifs in intervals that are uniquely bound in *D. melanogaster*. Again, this difference in motif conservation is statistically significant (Wilcoxon rank sum test, p = 6.04e-28). Taken together, these results show that, while DamID intervals that are bound by Dichaete *in vivo* in multiple species are not more highly conserved on average than those that are only bound in one species, individual Sox motifs within those intervals are both more likely to retain orthologous positions and to accumulate fewer mutations during evolution. Since DamID binding intervals are generally wider than ChIP-seq intervals and are not necessarily centred around the true binding site, it can be more difficult to identify bound motifs in DamID intervals than in ChIP intervals [95]. However, our findings highlight a link between *in vivo* Dichaete binding conservation and motif conservation, suggesting both functional importance of motif density and quality as well as a mechanism by which balancing selection at the sequence level could feed back to maintain functional binding events.

### High conservation of common binding by Dichaete and SoxN

Having observed that Dichaete binding shows a higher rate of conservation at functional sites such as known enhancers and core binding intervals, we asked whether a comparative approach could yield insights into the importance of common binding by Dichaete and SoxN. Previous studies have shown that Dichaete and SoxN show extensive binding similarity across the *D. melanogaster* genome and that in some cases they can compensate for each other’s loss at the level of DNA binding [51]. However, it is not known if widespread common binding has been retained throughout *Drosophila* evolution or its functional importance relative to unique binding by each TF. To address these questions, we turned to the Dichaete-Dam and SoxN-Dam data generated in *D. melanogaster* and *D. simulans*, first taking a qualitative approach to assess common and unique binding. Extensive common binding by Dichaete and SoxN was observed in both species, although, somewhat surprisingly, it represented a greater proportion of all binding events in *D. melanogaster* than in *D. simulans* (Figure 6A). Nonetheless, the single largest category of binding intervals we identified were those commonly bound by Dichaete and SoxN in both species (7415 intervals).

**Figure 6.**
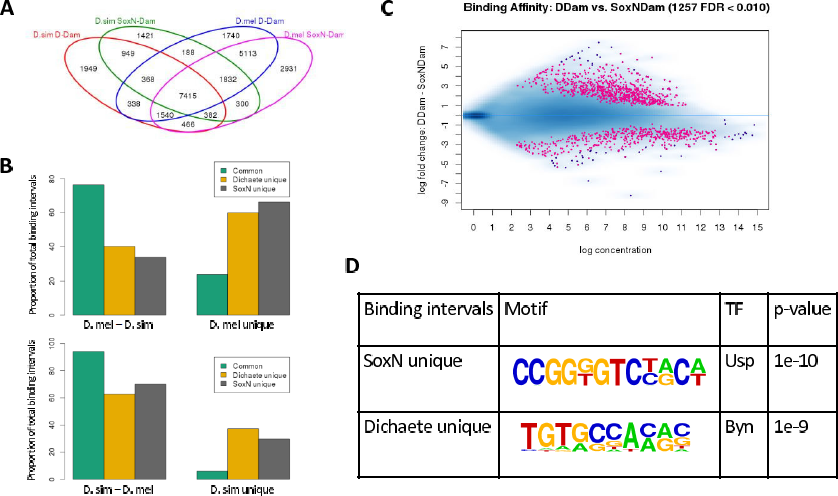
Preferential conservation of common binding by Dichaete and SoxN. (A). Venn diagram showing all Dichaete and SoxN binding intervals identified in *D. melanogaster* and *D. simulans*. The single largest category (7415 intervals) consists of commonly bound, conserved binding intervals. (B). Bar plots showing that commonly bound intervals are more likely to be conserved between *D. melanogaster* and *D. simulans* than intervals uniquely bound by either Dichaete or SoxN. The difference in conservation rates is highly significant for both *D. melanogaster* intervals (top, p < 2.2e-16 [approaches 0]) and *D. simulans* intervals (bottom, p < 2.2e-16 [approaches 0]). (C) MA plot showing binding intervals that are differentially bound between Dichaete (pink, fold change > 0) and SoxN (pink, fold change < 0) in both *D. melanogaster* and *D. simulans*. Fewer quantitative differences in binding are detected between the two TFs than between the binding profiles of one TF in two species. (D) Enriched motifs identified in intervals that are uniquely bound by SoxN or Dichaete in both *D. melanogaster* and *D. simulans*. A motif matching Ultraspiracle (Usp), a TF involved in axon pathfinding, was found in unique SoxN intervals, while a motif matching Brachyenteron (Byn), a TF critical for hindgut development, was found in unique Dichaete intervals.

Not only were there a large number of binding intervals commonly bound in both species, intervals that are commonly bound in either species are significantly more likely to be conserved than intervals bound uniquely by one Sox protein (Figure 6B). Out of all the intervals identified in *D. melanogaster* that are commonly bound by Dichaete and SoxN, 76.5% show binding conservation in *D. simulans*. In contrast, only 40.1% of uniquely bound Dichaete intervals are conserved in *D. simulans*, and 33.7% of uniquely bound SoxN intervals are conserved (Table 4), a highly significant difference between common and unique binding (χ^2^ = 3398.3, d.f. = 2, p < 2.2e-16 [approaches 0]). Performing the same analysis on the binding intervals identified in *D. simulans* reveals an even more striking effect, with 94.1% of commonly bound intervals showing binding conservation in *D. melanogaster*. Again, the difference in conservation rates between commonly and uniquely bound intervals is highly significant (χ^2^ = 2488.9, d.f. = 2, p < 2.2e-16 [approaches 0]). This effect is stronger than any of the previously examined functional categories, including known enhancers and core intervals.

**Table 4.**
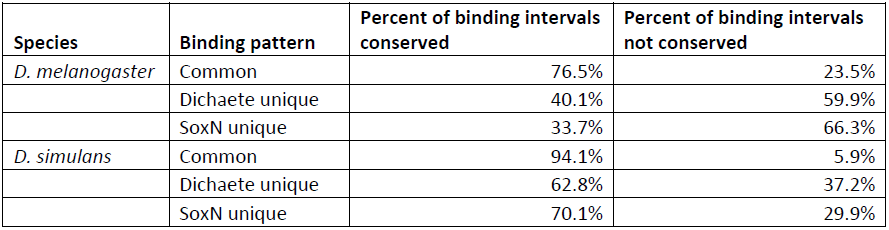
Conservation of common and unique binding by Dichaete and SoxN.

| Species | Binding pattern | Percent of binding intervals conserved | Percent of binding intervals not conserved |
| --- | --- | --- | --- |
| *D. melanogaster* | Common | 76.5% | 23.5% |
|  | Dichaete unique | 40.1% | 59.9% |
|  | SoxN unique | 33.7% | 66.3% |
| *D. simulans* | Common | 94.1% | 5.9% |
|  | Dichaete unique | 62.8% | 37.2% |
|  | SoxN unique | 70.1% | 29.9% |

Assigning these conserved, commonly bound intervals to putative target genes resulted in the identification of 5966 conserved core group B Sox targets in *D. melanogaster* and *D. simulans* (Additional File 4). These genes have a profile that is consistent with the known picture of group B Sox function. They are primarily upregulated in the developing CNS and are enriched for GO:BP terms related to biological regulation (p = 1.48e-49), system development (p = 2.86e-37), generation of neurons (p = 4.55e-31) and neuron differentiation (p = 4.92e-28) (Additional File 5). This suggests that many of the core functions of Dichaete and SoxN in the fly are achieved through regulation of genes at which both proteins can and do bind, and that this common, potentially redundant binding is evolutionarily conserved. To examine the conservation of these core group B targets on an expanded evolutionary scale, we compared them with the targets of the mouse group B Sox proteins Sox2 and Sox3 and the group C Sox protein Sox11 (Table 5). We mapped the targets of each of these proteins in neural precursor cells (NPCs, [29] to their *D. melanogaster* orthologues, resulting in a list of 1301 orthologues of Sox2 targets, 4213 orthologues of Sox3 targets and 1485 orthologues of Sox11 targets. Between 40-45% of the targets of each mouse Sox protein have conserved orthologues that are commonly bound by Dichaete and SoxN in the *Drosophila* genome, which is consistent with similar comparisons previously made between mouse Sox2 targets and targets of Dichaete and SoxN separately [51, 67]. Interestingly, roughly twice as many orthologues were found to be shared between Sox11 and SoxN alone as between Sox11 and the common targets of Dichaete and SoxN. This supports our previous suggestion that, while Dichaete and SoxN may contribute equally to homologous mammalian group B1 Sox functions, SoxN has evolved to take on a large part of the neural differentiation functions attributed to Sox11 independently of Dichaete. Overall, there is a high overlap between targets of mouse group B and C Sox proteins in the mammalian CNS and common, conserved targets of Dichaete and SoxN in the fly, supporting the deep evolutionary conservation of Sox functions in the CNS.

**Table 5.** Overlap between targets of mouse Sox proteins and common or core Dichaete and SoxN targets.

| Mouse protein | Orthologues of mouse targets in *D. melanogaster* | Overlap with common Dichaete/SoxN targets | Overlap with core SoxN targets | Overlap with core Dichaete targets |
| --- | --- | --- | --- | --- |
| Sox2 | 1301 | 589 | 443 | 522 |
| Sox3 | 4213 | 1730 | 1134 | 1590 |
| Sox11 | 1485 | 595 | 1092 | 610 |

While Dichaete and SoxN commonly bind the majority of conserved binding intervals, we also identified intervals that are uniquely bound by each TF in both species. We used DiffBind [86] to analyse quantitative differences in Dichaete and SoxN binding in both *D. melanogaster* and *D. simulans*. We detected 1257 binding intervals that are differentially bound between Dichaete and SoxN in both species (FDR1), with a greater number of intervals being preferentially bound by Dichaete (778) than SoxN (479) (Figure 6C). This is a considerably smaller difference than was seen when comparing Dichaete or SoxN binding between species, meaning that the binding profiles of these two paralogous TFs are less differentiated than the binding profiles of either Dichaete or SoxN in the genomes of two different species. We assigned both the preferentially bound intervals and the qualitatively uniquely bound intervals to target genes. This resulted in a set of 925 preferential Dichaete targets and 526 preferential SoxN targets, with 54 overlapping genes, and a set of 381 unique Dichaete targets and 361 unique SoxN targets, with only 14 overlapping genes (Additional Files 6 and 7).

Although the preferential and unique targets of each TF are not identical, they share some properties that can shed light on the unique functions of each protein. In both cases, while the two sets of genes have similar enrichments in terms of GO:BP terms, including terms related to morphogenesis, development, neuron differentiation and biological regulation (Additional Files 8 and 9), their spatial expression patterns according to FlyAtlas show different profiles. Both unique and preferential targets of SoxN are primarily upregulated in the larval CNS. The SoxN preferential target genes are enriched in the Reactome pathway *Role of Abl in Robo-Slit* signalling, an important pathway in axon guidance. Additionally, a *de novo* motif search in the uniquely bound SoxN intervals uncovered a motif corresponding to Ultraspiracle (Usp), a TF involved in several aspects of neuron morphogenesis [96, 97]. SoxN has been shown to be involved in the later aspects of CNS development, including axon pathfinding [51, 67], and its direct targets show high overlap with orthologous targets of mouse Sox11, which is active in post-mitotic, differentiating neurons [29]. Our results suggest that these functions may define the primary unique role of SoxN. On the other hand, Dichaete preferential and unique targets are upregulated in a wider range of tissues, including the brain, head, larval CNS, crop, eye, hindgut and thoracicoabdominal ganglion. Dichaete has previously been shown to be expressed and play a regulatory role in both the embryonic hindgut and brain [63, 67]. One of the top *de novo* motifs identified in the uniquely bound Dichaete intervals corresponds to Brachyenteron (Byn), a TF that is critical for the development of the hindgut [98, 99]. These results suggest that, while Dichaete and SoxN have many similar functions during development, key differences in their roles and target genes may be determined by differences in their own expression patterns, a signature of neofunctionalisation.

Our analysis of the genome-wide *in vivo* binding patterns of two group B Sox proteins in a comparative evolutionary context demonstrates a high degree of conservation in group B Sox function in the *Drosophila* species studied while identifying numerous instances of binding site turnover. In agreement with studies of other TFs in *Drosophila*, the amount of Dichaete binding site turnover between species and quantitative divergence in binding strength is correlated with phylogenetic distance [76, 77, 79]. However, we have also identified functional categories of sites which display significantly higher conservation than the background level, including binding intervals in known CRMs, binding intervals that have previously been identified as core intervals and intervals where both Dichaete and SoxN bind together. A two-way analysis of Dichaete and SoxN binding in two species of *Drosophila* reveals that the majority of conserved intervals are commonly bound by both TFs, leading us to identify a list of core group B Sox targets, while an analysis of uniquely bound conserved intervals provides indications as to the features that differentiate Dichaete and SoxN functions during development. Our results emphasize the fact that common, potentially redundant binding by Dichaete and SoxN has been specifically conserved during *Drosophila* evolution.

## Discussion

In this study, we have expanded upon our previous work identifying binding targets of Dichaete and SoxN in *D. melanogaster* to examine the evolutionary dynamics of group B Sox binding patterns in multiple species of *Drosophila*. Our analysis of Dichaete binding in four species showed high overall conservation in terms of target genes and genomic features associated with binding, but also uncovered extensive turnover in individual binding events. At the sequence level, we identified highly enriched Sox motifs in all sets of binding intervals that showed both density and nucleotide conservation rates correlated with binding conservation. To address the widespread common binding observed between Dichaete and SoxN, we performed a two-way comparison of the binding patterns of both TFs in two species, *D. melanogaster* and *D. simulans*. We observed that common binding is preferentially conserved between species in comparison to binding by a single factor alone, suggesting that Dichaete and SoxN’s ability to bind to the same loci is an important feature of their developmental functions. In addition, a large number of commonly bound targets are conserved, orthologous targets of group B Sox proteins in the mouse, particularly Sox2 and Sox3, which also show co-expression and functional redundancy in the developing CNS, raising the possibility of a deeply-conserved role for compensation among Sox group co-members.

We found a number of patterns in our three-way evolutionary analysis of Dichaete binding in *D. melanogaster*, *D. simulans* and *D. yakuba*. Normalizing the read counts from all samples together allowed us to reduce the effects of comparing separately thresholded datasets, which can lead to an underestimate of similarity. When we counted the divergent binding intervals between species, we found that the number of differences increases with the phylogenetic distance of the species being compared. A similar pattern was found in the correlations between binding profiles across all bound regions, with samples from more closely related species having higher correlations. These correlations, ranging from 0.62 – 0.72, are in line with the correlations between AP factor binding profiles in *D. melanogaster* and *D. yakuba*, which range from 0.57 for Cad to 0.75 for Kr [76]. We also identified instances of binding site turnover at individual gene loci, which could potentially represent compensatory gains and losses maintaining target gene expression levels during evolution. This mode of regulatory evolution appears to be important for Dichaete, as we found that in all pairwise comparisons, the majority of binding intervals that were not positionally conserved in one species had at least one binding interval present at a different position within the same locus in the other species. Interestingly, very few of these turnover events happened in CRMs that also displayed turnover between species [83], suggesting fluidity in individual binding site locations within relatively stable blocks of regulatory DNA [100]. Nonetheless, binding intervals associated with annotated CRMs from FlyLight or REDFly display a higher rate of conservation than those located outside of CRMs altogether, which highlights their functional role.

If balancing selection has worked to maintain functional binding events during *Drosophila* evolution, then one would expect to find selective constraint on the nucleotide sequence within binding intervals. Indeed, given the design of our DamID experiments, in which we expressed fusion proteins containing the *D. melanogaster* group B Sox sequences in each species, any differences in binding should be attributable to differences in the genome sequence or nuclear environment of each species, rather than to the Sox proteins themselves. Previous comparative studies using ChIP-seq have found that overall nucleotide conservation is not significantly elevated in conserved binding intervals [76, 77], and our findings agree with this observation. However, we did find significant correlations between binding conservation and Sox motif content in intervals. Conserved binding intervals have more matches to Sox motifs on average than non-conserved intervals or randomly chosen control intervals, indicating that an increased motif density may contribute to functional binding by group B Sox proteins and be an important facet in binding conservation. Conserved binding intervals also contain more Sox motifs that are positionally conserved across species and that show complete nucleotide conservation than non-conserved intervals. Additionally, Sox motifs within conserved intervals have a higher percentage of conserved nucleotides across all four species than those in non-conserved intervals or randomly shuffled control motifs. While we cannot determine from these data whether the presence of high-quality Sox motifs causes functional binding or whether functional binding leads to selective pressure to maintain Sox motifs, our results suggest that a feedback loop might operate between these two conditions, leading to the observed correlation between highly conserved motif matches and *in vivo* binding conservation.

One of the most perplexing aspects of group B Sox biology in both insects and vertebrates is their extensive co-expression, apparent functional redundancy and widespread common binding patterns. While we have previously described broad common binding as well as specific examples of binding compensation between Dichaete and SoxN in *D. melanogaster*, the functional and evolutionary significance of these patterns have remained unclear. Here we have found a clear preferential conservation of commonly bound intervals between *D. melanogaster* and *D. simulans*. These two species of *Drosophila* are closely related, showing a high amount of synteny and a level of divergence equivalent to that between human and the rhesus macaque, as measured by substitutions per neutral site [101, 102]. In agreement with previous studies, the rates of binding divergence for both Dichaete and SoxN between *Drosophila* species are lower than those for TFs in vertebrate species at comparable evolutionary distances [20, 81]. Nonetheless, despite the overall high level of binding conservation, the increase in conservation at commonly bound sites compared to uniquely bound sites, with 94% of commonly-bound *D. simulans* sites being conserved in *D. melanogaster*, is striking and highly significant. We expect that a comparison of group B Sox binding patterns in more distantly related *Drosophila* species will reveal even greater differences.

Integrating *in vivo* binding data from two species allowed us to identify a set of orthologous binding intervals that are consistently bound by both Dichaete and SoxN across genomes and nuclear environments. Many of the target genes associated with these intervals are deeply conserved group B Sox targets. Comparing these core target genes with the targets of mouse Sox2, Sox3 and Sox11 in the CNS revealed that the common, conserved targets of Dichaete and SoxN show greater overlap with both Sox2 and Sox3 targets than either core SoxN or Dichaete targets alone. On the other hand, the targets of Sox11, a group C Sox protein, show greater overlap with core SoxN targets than with either Dichaete or common targets in the fly. Taken together, these data suggest that common binding is an important feature of group B Sox function and that the roles played by Dichaete and SoxN through common binding are representative of ancient roles for group B Sox proteins that are conserved from insects to vertebrates.

The reasons for maintaining two paralogous TFs with such highly overlapping binding patterns and many compensatory functions during evolution remain unclear. One hypothesis is that conserved common binding is necessary to provide robustness to regulatory networks during critical stages of early neural development [51, 73, 103, 104]. However, common, conserved targets of Dichaete and SoxN include both genes where the two TFs have been shown to demonstrate functional compensation, such as the homeodomain DV-patterning genes *intermediate neuroblasts defective (ind)* and *ventral nervous system defective (vnd)*, as well as genes where they show at least some opposite regulatory effects, such as the proneural genes *achaete (ac)* and *lethal of scute (l’sc)* or *prospero (pros)*, a TF involved in neuroblast differentiation [51, 66, 67]. In addition to directly regulating the expression of target genes, Sox proteins can also induce DNA bending, potentially altering the local chromatin environment and creating indirect effects on gene expression by bringing other regulatory factors together [105–107]; such an architectural role might be more easily performed by multiple family members than target-specific functions. These observations suggest a complex functional relationship between group B Sox family members in flies, including both functional compensation and balanced, opposing effects at certain loci, both of which are evolutionarily conserved. The high overlap in the regulatory networks influenced by Dichaete and SoxN [51] might itself lead to selective constraint on binding sites, as mutations affecting sites that can be functionally bound by both TFs would have a greater disruptive effect. This is consistent with the highly conserved DNA-binding domains found in paralogous group B Sox proteins.

A model of strong selection to maintain common binding between Dichaete and SoxN can also explain the types of neofunctionalisations that they have acquired. At the sequence level, the majority of the diversification between *Dichaete* and *SoxN* can be found in regions outside of the HMG domain, which could drive interactions with specific binding partners, and in each gene’s regulatory regions, which determine where they are uniquely or co-expressed [45]. Although they show extensive of overlapping expression in the developing CNS, Dichaete and SoxN are also expressed in specific domains in the embryo; Dichaete shows unique expression in the midline, brain and hindgut, while SoxN is expressed uniquely in the lateral column of the neuroectoderm and in a specific pattern in the epidermis at later stages [50, 55, 63, 108]. The functional signatures of targets genes that we found to be uniquely bound and evolutionarily conserved, including both spatial expression patterns and enriched motifs corresponding to potential cofactors, point to these domain-specific roles. Although changes in target specificity due to binding with cofactors has not been demonstrated for Sox proteins in *Drosophila*, it is a well-characterized phenomenon for the Hox family of TFs [11, 109, 110]. Dichaete has previously been shown to bind together with Ventral veins lacking (Vvl) in the midline [50, 56]; we have identified an enriched motif for the hindgut-specific factor Byn in uniquely bound, conserved Dichaete intervals, which may represent a similar interaction.

Unlike vertebrate group B Sox proteins, which can be split into two subgroups with largely specialised regulatory functions, Dichaete and SoxN appear to act as partially redundant activators at a large set of core target genes and to oppose each other’s activity at a smaller set of targets. In addition, each protein uniquely regulates smaller subsets of genes, both positively and negatively, in specific tissues [51]. These differences reflect the independent evolutionary trajectories of group B Sox genes in vertebrates and insects, which are still not fully resolved [45, 60]. However, given the broad patterns of co-expression and functional redundancy observed between co-members of various Sox families across the Sox phylogeny [27, 31, 32, 37–43, 49], it is logical to ask whether such partial redundancy is a shared ancestral feature or the result of convergent evolution. Two competing models for the evolution of group B *Sox* genes both propose at least one duplication event at the ancestral *SoxB* locus before the protostome/deuterostome split, either in tandem or as the result of a whole genome duplication [60]. While the details of the group B *Sox* expansion are debated, it is possible that redundancy between the first group B *Sox* paralogues may have been retained throughout evolution, while later paralogues split into group B1 and B2 in vertebrates or acquired partial neofunctionalisations in insects. This model, combined with our data, suggests that the integrated action of multiple Sox proteins is an ancestral feature of CNS development that has been consistently selected for and that continues to be elaborated on and refined in different lineages.

## Conclusions

The studies presented here have examined the *in vivo* binding of the *Drosophila* group B Sox proteins Dichaete and SoxN in an evolutionary context, focusing on a detailed analysis of the patterns of binding site turnover for Dichaete in four species and a multi-factor analysis of Dichaete and SoxN binding in two species. We show broad conservation of Dichaete and SoxN regulatory networks coupled with widespread binding divergence at a rate that is consistent with studies of other TFs in *Drosophila*. We demonstrate preferential binding conservation at functional sites, including known CRMs, as well as even stronger binding conservation at sites that are commonly bound by Dichaete and SoxN. The common, conserved binding intervals define a set of targets that also share deep orthology with targets of mouse group B Sox proteins, suggesting that they represent ancestral functions in the CNS. Finally, we propose a mode of group B Sox evolution whereby common binding and partial redundancy is specifically maintained, while individual paralogues acquire novel functions largely through changes to their own expression patterns and/or binding partners.

## Methods

### Fly husbandry and embryo collection

The wild-type strains of the following *Drosophila* species were used in all experiments: *D. melanogaster w^1118^; D. simulans w^[501]^* (reference strain - http://www.ncbi.nlm.nih.gov/genome/200?genome_assembly_id=28534); *D. yakuba Cam-115* [111]; *D. pseudoobscura pseudoobscura* (reference strain - http://www.ncbi.nlm.nih.gov/genome/219?genome_assembly_id=28567). *D. melanogaster*, *D. simulans* and *D. yakuba* stocks were kept at 25° C on standard cornmeal medium. *D. pseudoobscura* stocks were kept at 22.5° C at low humidity on banana-opuntia-malt medium (1000 ml water, 30 g yeast, 10 g agar, 20 ml Nipagin, 150 g mashed bananas, 50 g molasses, 30 g malt, 2.5 g opuntia powder). All embryo collections were performed at 25° C on grape juice agar plates supplemented with fresh yeast paste. Embryos were dechorionated for 3 minutes in 50% bleach and washed with water before snap-freezing in liquid nitrogen and storage at K80° C.

### Generation of transgenic lines

Three piggyBac vectors were created for DamID, containing a Dichaete-Dam construct, a SoxN-Dam construct or a Dam-only construct. The SoxN-Dam fusion protein coding sequence, along with upstream UAS sites, an *Hsp70* promoter and the *SV40 5'* UTR, was cloned from an existing pUAST vector [51]. The Dichaete-Dam fusion protein coding sequence, along with upstream UAS sites, an *Hsp70* promoter and the *kayak* 5’ UTR, was cloned from genomic DNA extracted from a *D. melanogaster* line carrying this construct [112]. The Dam coding region, as well as upstream UAS sites, an *Hsp70* promoter and the *SV40* 5’ UTR, was directly excised from an existing pUAST vector (gift from T. Southall) using *Sph*I and *Stu*I. All inserts were first cloned into the pSLfa1180fa shuttle vector (gift from E. Wimmer) [113] (SoxN-Dam: *Spe*I/*Stu*I, Dichaete-Dam: *Spe*I/*Avr*II, Dam: *Sph*I/*Stu*I). The inserts were then excised from the shuttle vectors using the octoKcutters *Fse*I and *Asc*I and cloned into the final pBac3xP3-EGFP vector (also from Ernst Wimmer). Plasmid DNA was microinjected into embryos from each species at a concentration of 0.6 μg/μl, together with a piggyBac helper plasmid at 0.4 μg/μl. All microinjections were performed by Sang Chan in the Department of Genetics injection facility, University of Cambridge. Surviving adults were backcrossed to *w; Sco/SM6a* males or virgin females for *D. melanogaster* or to males or virgin females from the parental line for the other species. F1 progeny were scored for eyeKspecific GFP expression, and *D. melanogaster* insertions were balanced over either *SM6a* or *TM6c*.

### Isolation of DamID DNA and sequencing

Embryos from the *Dichaete-Dam*, *SoxN-Dam* and *Dam-only* transgenic lines in each species were collected after overnight lays and DNA was isolated using minor modifications to the protocol of Vogel and colleagues [95]. Three biological replicates were collected from each line, consisting of approximately 50-150 μl settled volume of embryos per replicate. To extract high-molecular weight genomic DNA, each aliquot of embryos was homogenized in a Dounce 15-ml homogenizer in 10 ml of homogenization buffer. 10 strokes were applied with pestle B, followed by 10 strokes with pestle A. The lysate was then spun for 10 minutes at 6000g. The supernatant was discarded, and the pellet was resuspended in 10 ml homogenization buffer, then spun again for 10 minutes at 6000g. The supernatant was again discarded, and the pellet was resuspended in 3 ml homogenization buffer. 300 μl of 20% n-lauroyl sarcosine were added, and the samples were inverted several times to lyse the nuclei. The samples were treated with RNaseA followed by proteinase K at 37° C. They were then purified by two phenol-chloroform extractions and one chloroform extraction. Genomic DNA was precipitated by adding 2 volumes of EtOH and 0.1 volume of 3M NaOAc, dried and resuspended in 50-150 μl TE buffer, depending on the starting amount of embryos.

30 μl of each DNA sample was digested for at least 2 hours with *Dpn*I at 37° C. To eliminate observed contamination from non-digested genomic DNA, 0.7 volumes of Agencourt AMPure XP beads (Beckman Coulter) were added to each sample to remove high-molecular weight DNA. 1.1 volumes of Agencourt AMPure XP beads were then added to recover all remaining DNA fragments, which were eluted in 30 μl TE buffer and then ligated to a double-stranded oligonucleotide adapter. If bands were observed in the PCR products, the ligation was repeated with adapters titrated to 1:2 or 1:4. Ligation products were digested with DpnII to remove fragments with unmethylated GATC sequences and amplified by PCR, followed by purification via a phenol-chloroform extraction. The DNA was precipitated and resuspended in 50 μl TE buffer, then sonicated in order to reduce the average fragment size using a Covaris S2 sonicator with the following settings: intensity 5, duty cycle 10%, 200 cycles/burst, 300 seconds. The samples were purified using a QIAquick PCR Purification kit to remove small fragments. Sample concentrations were measured using a Qubit with the DNA High Sensitivity Assay (Life Technologies), and the size distributions of DNA fragments were measured using a 2100 Bioanalyzer with the High Sensitivity DNA kit and chips (Agilent). Samples were sent to BGI Tech Solutions (HongKong) Co., Ltd., for library construction using the standard ChIP-seq library protocol and were sequenced on an Illumina MiSeq or HiSeq 2000. Libraries were multiplexed with 2 samples per run for the MiSeq and 9-12 samples per lane for the HiSeq. MiSeq libraries were run as 150-bp single-end reads, while HiSeq libraries were run as 50-bp single-end reads.

### Sequencing data analysis

Cutadapt [114] was used to trim DamID adapter sequences from both ends of reads. Trimmed reads were mapped against the following reference genomes using bowtie2 [115] with the default settings: *D. melanogaster* April 2006 (UCSC dm3, BDGP) [116, 117], *D. simulans* April 2005 (UCSC droSim1, The Genome Institute at Washington University (WUSTL)) [101], *D. yakuba* November 2005 (UCSC droYak2, The Genome Institute at WUSTL) [101] and *D. pseudoobscura* November 2004 (UCSC dp3, Baylor College of Medicine Human Genome Sequencing Center (BCM-HGSC)) [101, 118]. All reference genomes were downloaded from the UCSC Genome Browser (http://hgdownload.soe.ucsc.edu/downloads.html). Mapped reads were sorted and indexed using SAMtools [119].

For DamID data analysis, the position of every GATC site in each genome was determined using the HOMER utility scanMotifGenomeWide.pl (http://homer.salk.edu/homer/index.html) [120]. For each sample, reads were extended to the average fragment length (200 bp) using the BEDTools slop utility. The number of extended reads overlapping each GATC fragment was then calculated using the BEDTools coverage utility [90]. The resulting counts for each sample were collated to form a count table, consisting of one column for each fusion protein or Dam-only sample and one row for each GATC fragment. The count tables served as inputs to run DESeq2 (run in R version 3.1.0 using RStudio version 0.98), which was used to test for differential enrichment in the fusion protein samples versus the Dam-only samples at each GATC fragment [121]. Fragments flagged as differentially enriched (log2 fold change > 0 and adjusted p-value < 0.05 or < 0.01) were extracted, and neighbouring enriched fragments within 100 bp were merged to formed binding intervals. Binding intervals were scanned for *de novo* and known motifs using HOMER findMotifsGenome.pl [120]. An overview of this pipeline can be found at https://github.com/sarahhcarl/flychip/wiki/Basic-DamlD-analysis-pipeline.

Both binding intervals and reads from non-*D. melanogaster* species were translated to the *D. melanogaster* UCSC dm3 reference genome using the LiftOver utility from the UCSC Genome Browser (http://genome.ucsc.edu/cgi-bin/hgLiftOver) [122]. For *D. simulans* and *D. yakuba*, the minMatch parameter was set to 0.7, while for *D. pseudoobscura* it was set to 0.5; multiple outputs were not permitted. DiffBind was used to enable a quantitative comparison between DamID datasets from all species using translated data [86]. Translated reads in BED format were back-converted to SAM format using a custom perl script (bed2sam.pl, https://github.com/sarahhcarl/Flychip/tree/master/DamID_analysis), and then into BAM format using SAMtools [119]. For each analysis, the translated reads from each sample were normalized together in DiffBind using the DESeq2 normalization method.

Binding intervals were assigned to the closest gene in the *D. melanogaster* genome using a perl script written by Bettina Fischer (University of Cambridge). Genomic feature annotations were performed using ChIPSeeker [123]. The distances from binding intervals to TSSs were calculated and plotted using ChIPpeakAnno [124]. All calculations of overlaps between interval datasets were performed using the BEDTools intersect utility [90]. All visualization of sequence data was done using the Integrated Genome Browser (IGB) [125]. Gene ontology analysis was performed using FlyMine [126], with a Benjamini and Hochberg correction for multiple testing and a correction for gene length effect.

### Evolutionary sequence analysis of binding motifs

DiffBind was used to identify the set of Dichaete-Dam binding intervals unique to *D. melanogaster* and the set conserved in all four species [86]. The genomic coordinates of these intervals in *D. melanogaster* were extracted and translated to each other species’ reference genome assembly using LiftOver, with the minMatch parameter set to 0.7. The sequences of each orthologous binding interval in each species were obtained using the fetch-UCSC sequences tool from RSAT, preserving strand information [93]. For each interval for which one unambiguous orthologous sequence could be identified in all four species, the sequences were multiply aligned using PRANK, estimating the guide tree directly from the data [91, 92]. Two strategies were used to predict Sox binding sites within binding intervals. First, the positional weight matrices (PWMs) representing the top-scoring *de novo* Sox motif identified via HOMER in each binding interval dataset were downloaded [120]. FIMO was used to search for matches to each PWM in all binding intervals, and the resulting hits were used to calculate the average number of motifs per binding interval [89]. The PWMs were also used to scan multiple alignments of both four-way conserved and unique *D. melanogaster* Dichaete-Dam binding intervals using matrix-scan from RSAT, with the pre-compiled *Drosophila* background file and with the cutoff for reporting matches set to a PWM weight-score of >= 4 and a p-value of < 0.0001. If a binding interval was identified at the same aligned position in an orthologous binding interval in more than one species, it was considered to be positionally conserved between those species. Randomly shuffled control motifs were generated using the RSAT tool permute-matrix [93] and used in the same way. All statistical tests were performed in R v.3.1.0 using RStudio v0.98.

### Data access

All sequencing and binding interval data described in this paper have been deposited in NCBI’s Gene Expression Omnibus (GEO) [127] and are accessible through GEO Series accession number GSE63333 (http://www.ncbi.nlm.nih.gov/geo/query/acc.cgi?acc=GSE63333).

## Competing interests

The authors declare that they have no competing interests.

### Authors’ contributions

SC and SR conceived and designed the experiments; SC performed the experiments and analysed the data; SC and SR drafted the manuscript. All authors read and approved the final manuscript.

### Description of additional files

**Supplementary Figure 1.**
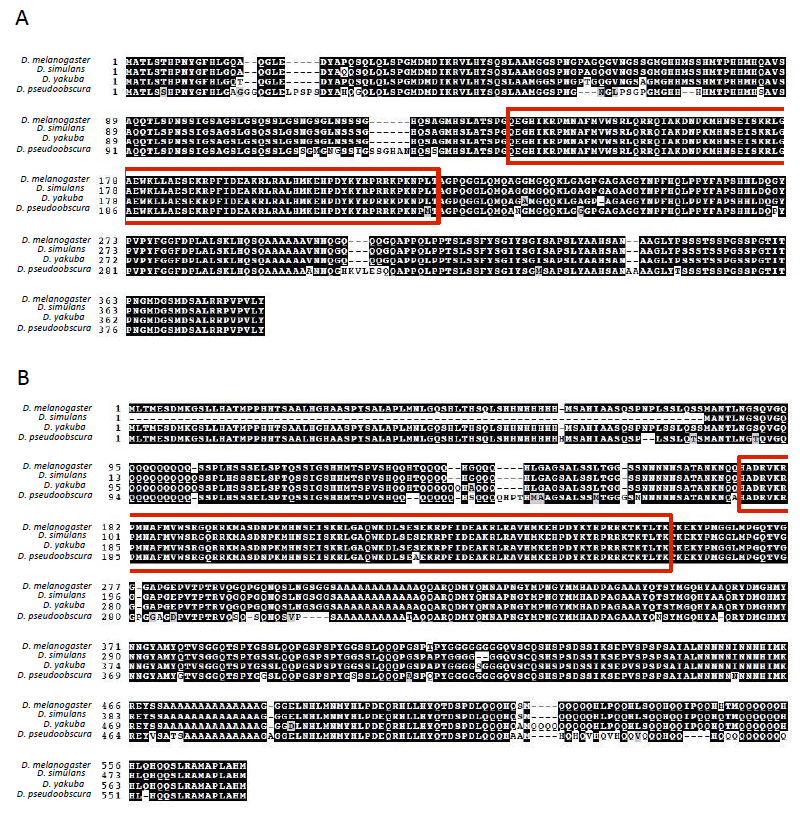
Multiple alignment of Dichaete and SoxNeuro amino acid sequences. (A) Multiple alignment of the entire Dichaete sequence from *D. melanogaster, D. simulans, D. yakuba* and *D. pseudoobscura*. (B) Multiple alignment of the entire SoxN sequence from *D. melanogaster*, *D. simulans*, *D. yakuba* and *D. pseudoobscura*. The HMG domains of each orthologous protein are highlighted in red.

**Supplementary Figure 2.**
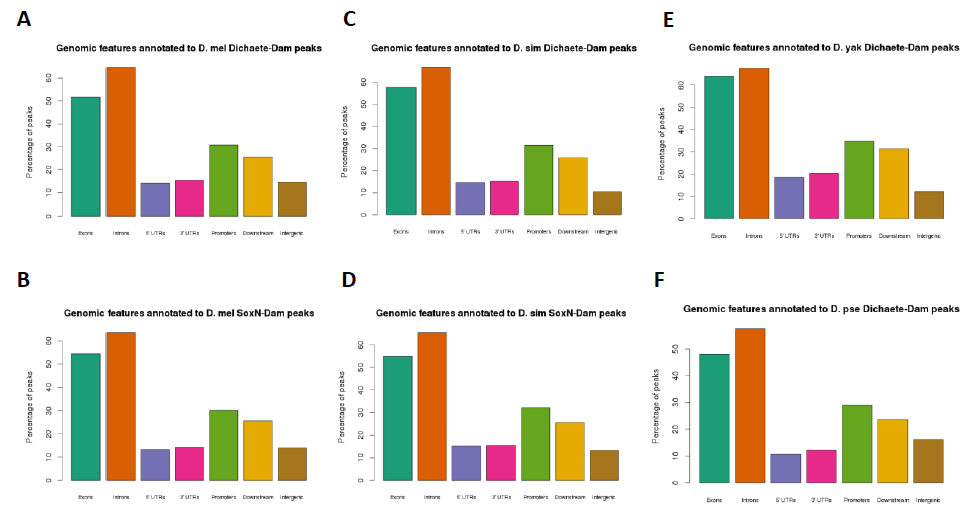
Genomic features annotated to group B Sox DamID binding intervals. Feature classes include exons, introns, 5’ UTRs, 3’UTRs, promoters, immediate downstream and intergenic. Each interval may be annotated with more than one class if it overlaps multiple features. (A) Percentages of *D. melanogaster* Dichaete-Dam intervals annotated to each feature class. (B) Percentages of *D. melanogaster* SoxN-Dam intervals annotated to each feature class. (C) Percentages of *D. simulans* Dichaete-Dam intervals annotated to each feature class. (D) Percentages of *D. simulans* SoxN-Dam intervals annotated to each feature class. (E) Percentages of *D. yakuba* Dichaete-Dam intervals annotated to each feature class. (F) Percentages of *D. pseudoobscura* Dichaete-Dam intervals annotated to each feature class.

**Supplementary Figure 3.**
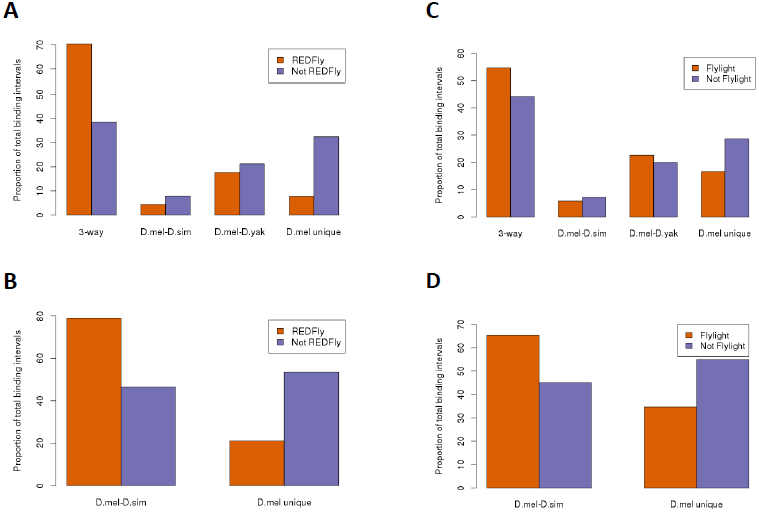
DamID intervals overlapping a known CRM are preferentially conserved. (A) Dichaete-Dam binding intervals that overlap a REDFly CRM are more likely to show three-way binding conservation between *D. melanogaster*, *D. simulans* and *D. yakuba* (“3-way”) and are less likely to be unique to *D. melanogaster* (“D. mel unique”) than those that do not (p = 7.O6e-35). (B) SoxN-Dam binding intervals that overlap a REDFly CRM are more likely to show two-way binding conservation between *D. melanogaste* and *D. simulans* (“D.mel-D.sim”) and are less likely to be unique to *D. melanogaste* than those that do not (p = 2.4Oe-72). (C) Dichaete-Dam binding intervals that overlap a FlyLight CRM are more likely to show three-way binding conservation and are less likely to be unique to *D. melanogaste* than those that do not (p = 3.38e-38). (D) SoxN-Dam binding intervals that overlap a FlyLight CRM are more likely to show two-way binding conservation and are less likely to be unique to *D. melanogaste* than those that do not (p = 7.27e-12).

**Supplementary Figure 4.**
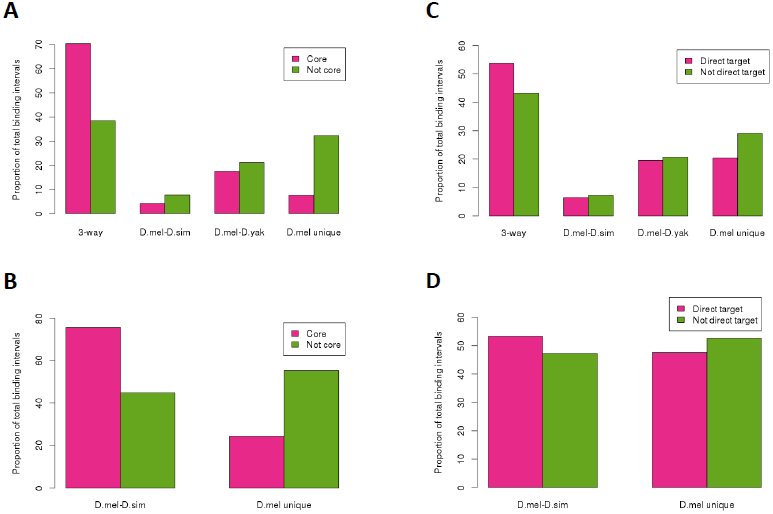
DamID intervals overlapping a core binding interval or annotated to a direct target gene are preferentially conserved. (A) Dichate-Dam binding intervals that overlap a core Dichaete binding site are more likely to show three-way conservation between *D. melanogaster*, *D. simulans* and *D. yakuba* (“3-way”) and are less likely to be unique to *D. melanogaste* (“D. mel unique”) than those that do not (p = 4.10e-305). (B) SoxN-Dam binding intervals that overlap a core SoxN binding site are more likely to show two-way conservation between *D. melanogaste* and *D. simulans* (“2-way”) and are less likely to be unique to *D. melanogaste* than those that do not (p = 1.90e-161). (C) Dichaete-Dam binding intervals that are annotated to a Dichaete direct target gene are more likely to show three-way conservation and are less likely to be unique to *D. melanogaste* than those that are not (p = 2.3e-12). However, this effect is weaker than for core intervals. (D) SoxN-Dam binding intervals that are annotated to a SoxN direct target gene are more likely to show two-way conservation and are less likely to be unique to *D. melanogaste* than those that are not (p = 3.4e-5). Again, however, this effect is weaker than for core intervals.

## Additional file 1

File format: .xls

Title: Genes annotated to Sox DamID binding intervals

Description: List of target genes annotated to Dichaete and SoxN DamID binding intervals in all species. For *non-melanogaster* species, the annotation was performed on binding intervals called from sequence data that had been translated to the *D. melanogaste* genome. The CG numbers from NCBI, gene symbols and Flybase identifiers for each gene are listed.

### Additional file 2

File format: .xls

Title: Gene ontology terms enriched in Sox target gene lists

Description: List of all significantly enriched GO terms for Dichaete and SoxN annotated target genes in all species. The first column lists the GO biological process term and the second column lists the corrected p-value.

### Additional file 3

File format: .xls

Title: Enriched *de novo* motifs discovered in Sox binding intervals.

Description: For each Dichaete and SoxN DamID binding interval dataset, the top 20 enriched *de novo* motifs discovered by HOMER are listed. The first column lists the rank of the motif, the second column lists the best guess TF matching the motif according to HOMER, the third column lists the consensus sequence of the motif and the fourth column lists its p-value.

### Additional file 4

File format: .xls

Title: Target genes annotated to conserved, commonly bound intervals.

Description: List of target genes annotated to binding intervals that are commonly bound by Dichaete and SoxN as well as conserved between *D. melanogaste* and *D. simulans*. The CG numbers from NCBI, gene symbols and Flybase identifiers for each gene are listed.

### Additional file 5

File format: .xls

Title: Gene ontology terms enriched in commonly bound, conserved target genes.

Description: List of all significantly enriched GO terms for target genes annotated to intervals that are commonly bound by Dichaete and SoxN and are conserved between *D. melanogaste* and *D. simulans*. The first column lists the GO biological process term and the second column lists the corrected p-value.

### Additional file 6

File format: .xls

Title: Target genes annotated to conserved, uniquely bound Dichaete intervals.

Description: List of target genes annotated to binding intervals that are uniquely bound by Dichaete and conserved between *D. melanogaste* and *D. simulans*. The CG numbers from NCBI, gene symbols and Flybase identifiers for each gene are listed.

### Additional file 7

File format: .xls

Title: Gene ontology terms enriched in conserved, uniquely bound Dichaete target genes.

Description: List of all significantly enriched GO terms for target genes annotated to intervals that are uniquely bound by Dichaete and are conserved between *D. melanogaste* and *D. simulans*. The first column lists the GO biological process term and the second column lists the p-value.

### Additional file 8

File format: .xls

Title: Target genes annotated to conserved, uniquely bound SoxN intervals.

Description: List of target genes annotated to binding intervals that are uniquely bound by SoxN and conserved between *D. melanogaste* and *D. simulans*. The CG numbers from NCBI, gene symbols and Flybase identifiers for each gene are listed.

### Additional file 9

File format: .xls

Title: Gene ontology terms enriched in conserved, uniquely bound SoxN target genes.

Description: List of all significantly enriched GO terms for target genes annotated to intervals that are uniquely bound by SoxN and are conserved between *D. melanogaste* and *D. simulans*. The first column lists the GO biological process term and the second column lists the p-value.

## Acknowledgements

This work was supported by a Wellcome Trust 4-year Ph.D. studentship and a Cambridge Overseas Trust studentship to S.C. The funders had no role in study design, data collection and analysis, decision to publish, or preparation of the manuscript. We thank Tony Southall for providing the pUAST-Dam vector and Enrico Ferrero for providing the pUAST-SoxNDam vector. We also thank Ernst Wimmer for providing the pSLfa1180fa vector, the pBac3xP3-EGFP vector and the piggyBac transposase helper plasmid. We are indebted to Sang Chan for performing microinjections and to Bettina Fischer for support in the lab and insightful discussions about data analysis.

## References

1. Dunham I, Kundaje A, Aldred SF, Collins PJ, Davis CA, Doyle F, Epstein CB, Frietze S, Harrow J, Kaul R, Khatun J, Lajoie BR, Landt SG, Lee B-K, Pauli F, Rosenbloom KR, Sabo P, Safi A, Sanyal A, Shoresh N, Simon JM, Song L, Trinklein ND, Altshuler RC, Birney E,Brown JB, Cheng C, Djebali S, Dong X Dunham I, et al.: An integrated encyclopedia of DNA elements in the human genome. Nature 2012, 489:57–74.

2. Gordon KL, Ruvinsky I: Tempo and mode in evolution of transcriptional regulation. PLoS Genet 2012, 8:e1002432.

3. Neph S, Vierstra J, Stergachis AB, Reynolds AP, Haugen E, Vernot B, Thurman RE, John S, Sandstrom R, Johnson AK, Maurano MT, Humbert R, Rynes E, Wang H, Vong S, Lee K, Bates D, Diegel M, Roach V, Dunn D, Neri J, Schafer A, Hansen RS, Kutyavin T, Giste E, Weaver M, Canfield T, Sabo P, Zhang M, Balasundaram G, et al.: An expansive human regulatory lexicon encoded in transcription factor footprints. Nature 2012, 489:83–90.

4. The modENCODE Consortium, Roy S, Ernst J, Kharchenko PV, Kheradpour P, Negre N, Eaton ML, Landolin JM, Bristow CA, Ma L, Lin MF, Washietl S, Arshinoff BI, Ay F, Meyer PE, Robine N, Washington NL, Di Stefano L, Berezikov E, Brown CD, Candeias R, Carlson JW, Carr A, Jungreis I, Marbach D, Sealfon R, Tolstorukov MY, Will S, Alekseyenko AA, Artieri C, et al.: Identification of Functional Elements and Regulatory Circuits by Drosophila modENCODE. Science 2010, 330:1787–1797.

5. Wray GA: The evolutionary significance of cis-regulatory mutations. Nat Rev Genet 2007, 8:206–216.

6. Biggin MD: Animal Transcription Networks as Highly Connected, Quantitative Continua. Dev Cell 2011, 21:611–626.

7. Hueber SD, Lohmann I: Shaping segments: Hox gene function in the genomic age. BioEssays 30:965–979.

8. Hueber SD, Bezdan D, Henz SR, Blank M, Wu H, Lohmann I: Comparative analysis of Hox downstream genes in Drosophila. Development 2007, 134:381–392.

9. Kaplan T, Li X-Y, Sabo PJ, Thomas S, Stamatoyannopoulos JA, Biggin MD, Eisen MB: Quantitative Models of the Mechanisms That Control Genome-Wide Patterns of Transcription Factor Binding during Early Drosophila Development. PLoS Genet 2011, 7:e1001290.

10. Li X-Y, Thomas S, Sabo PJ, Eisen MB, Stamatoyannopoulos JA, Biggin MD: The role of chromatin accessibility in directing the widespread, overlapping patterns of Drosophila transcription factor binding. Genome Biol 2011, 12:R34.

11. Mann RS, Lelli KM, Joshi R: Chapter 3 Hox Specificity. In Curr Top Dev Biol. Volume 88. Elsevier; 2009:63–101.

12. Zinzen RP, Girardot C, Gagneur J, Braun M, Furlong EEM: Combinatorial binding predicts spatioV temporal cisVregulatory activity. Nature 2009, 462:65–70.

13. Aleksic J, Russell S: ChIPing away at the genome: the new frontier travel guide. Mol Biosyst 2009, 5:1421.

14. Chen Y, Negre N, Li Q, Mieczkowska JO, Slattery M, Liu T, Zhang Y, Kim T-K, He HH, Zieba J, Ruan Y, Bickel PJ, Myers RM, Wold BJ, White KP, Lieb JD, Liu XS: Systematic evaluation of factors influencing ChIP-seq fidelity. Nat Methods 2012, 9:609–614.

15. Fisher WW, Li JJ, Hammonds AS, Brown JB, Pfeiffer BD, Weiszmann R, MacArthur S, Thomas S, Stamatoyannopoulos JA, Eisen MB, Bickel PJ, Biggin MD, Celniker SE: DNA regions bound at low occupancy by transcription factors do not drive patterned reporter gene expression in Drosophila. Proc Natl Acad Sci 2012, 109:21330–21335.

16. Marinov GK, Kundaje A, Park PJ, Wold BJ: LargeVScale Quality Analysis of Published ChIP-seq Data. G358 GenesGenomesGenetics 2013, 4:209–223.

17. Park D, Lee Y, Bhupindersingh G, Iyer VR: Widespread Misinterpretable ChIP-seq Bias in Yeast. PloS One 2013, 8:e83506.

18. Kim J, He X, Sinha S: Evolution of Regulatory Sequences in 12 Drosophila Species. PLoS Genet 5:e1000330.

19. Ludwig MZ, Bergman C, Patel NH, Kreitman M: Evidence for stabilizing selection in a eukaryotic enhancer element. Nature 2000, 403:564–567.

20. Villar D, Flicek P, Odom DT: Evolution of transcription factor binding in metazoans — mechanisms and functional implications. Nat Rev Genet 2014.

21. Jager M, Queinnec E, Houliston E, Manuel M: Expansion of the SOX gene family predated the emergence of the Bilateria. Mol Phylogenet Evol 2006, 39:468–477.

22. Jager M, Queinnec E, Chiori R, Le Guyader H, Manuel M: Insights into the early evolution of SOX genes from expression analyses in a ctenophore. J Exp Zoolog B Mol Dev Evol 2008, 310B:650–667.

23. Larroux C, Fahey B, Liubicich D, Hinman VF, Gauthier M, Gongora M, Green K, Wörheide G, Leys SP, Degnan BM: Developmental expression of transcription factor genes in a demosponge: insights into the origin of metazoan multicellularity. Evol Dev 2006, 8:150–173.

24. Phochanukul N, Russell S: No backbone but lots of Sox: Invertebrate Sox genes. Int J Biochem Cell Biol 2010, 42:453–464.

25. Srivastava M, Begovic E, Chapman J, Putnam NH, Hellsten U, Kawashima T, Kuo A, Mitros T, Salamov A, Carpenter ML, Signorovitch AY, Moreno MA, Kamm K, Grimwood J, Schmutz J, Shapiro H, rigoriev IV, Buss LW, Schierwater B, Dellaporta SL, Rokhsar DS: The Trichoplax genome and the nature of placozoans. Nature 2008, 454:955–960.

26. Bowles J, Schepers G, Koopman P: Phylogeny of the SOX Family of Developmental Transcription Factors Based on Sequence and Structural Indicators. Dev Biol 2000, 227:239–255.

27. Guth SIE, Wegner M: Having it both ways: Sox protein function between conservation and innovation. Cell Mol Life Sci 2008, 65:3000–3018.

28. Schepers GE, Teasdale RD, Koopman P: Twenty pairs of sox: extent, homology, and nomenclature of the mouse and human sox transcription factor gene families. Dev Cell 2002: 167–170.

29. Bergsland M, Ramskold D, Zaouter C, Klum S, Sandberg R, Muhr J: Sequentially acting Sox transcription factors in neural lineage development. Genes Dev 2011, 25:2453–2464.

30. Kamachi Y, Uchikawa M, Collignon J, Lovell-Badge R, Kondoh H: Involvement of Sox1, 2 and 3 in the early and subsequent molecular events of lens induction. Development 1998, 125:2521–2532.

31. Uwanogho D, Rex M, Cartwright EJ, Pearl G, Healy C, Scotting PJ, Sharpe PT: Embryonic expression of the chicken Sox2, Sox3 and Sox11 genes suggests an interactive role in neuronal development. Mech Dev 1995, 49:23–36.

32. Wood HB, Episkopou V: Comparative expression of the mouse Sox1, Sox2 and Sox3 genes from preVgastrulation to early somite stages. Mech Dev 1999, 86:197–201.

33. Huang J, Arsenault M, Kann M, Lopez-Mendez C, Saleh M, Wadowska D, Taglienti M, Ho J, Miao Y, Sims D, Spears J, Lopez A, Wright G, Hartwig S: The transcription factor sryVrelated HMG boxV4 (SOX4) is required for normal renal development *in vivo*: *SOXC* Genes During Renal Development. Dev Dyn 2013, 242:790–799.

34. Sock E, Rettig SD, Enderich J, Bosl MR, Tamm ER, Wegner M: Gene Targeting Reveals a Widespread Role for the High-Mobility-Group Transcription Factor Sox11 in Tissue Remodeling. Mol Cell Biol 2004, 24:6635–6644.

35. Wilson ME, Yang KY, Kalousova A, Lau J, Kosaka Y, Lynn FC, Wang J, Mrejen C, Episkopou V, Clevers HC, German MS: The HMG Box Transcription Factor Sox4 Contributes to the Development of the Endocrine Pancreas. Diabetes 2005, 54:3402–3409.

36. Downes M, Koopman P: SOX18 and the Transcriptional Regulation of Blood Vessel Development. Trends Cardiovasc Med, 11:318–324.

37. Matsui T: Redundant roles of Sox17 and Sox18 in postnatal angiogenesis in mice. J Cell Sci 2006, 119:3513–3526.

38. Bhattaram P, Penzo-Méndez A, Sock E, Colmenares C, Kaneko KJ, Vassilev A, DePamphilis ML, Wegner M, Lefebvre V: Organogenesis relies on SoxC transcription factors for the survival of neural and mesenchymal progenitors. Nat Commun 2010, 1:1–12.

39. Ferri ALM: Sox2 deficiency causes neurodegeneration and impaired neurogenesis in the adult mouse brain. Development 2004, 131:3805–3819.

40. Nishiguchi S, Wood H, Kondoh H, Lovell-Badge R, Episkopou V: Soxl directly regulates the y-crystallin genes and is essential for lens development in mice. Genes Dev 1998, 12:776–781.

41. Okuda Y, Ogura E, Kondoh H, Kamachi Y: B1 SOX Coordinate Cell Specification with Patterning and Morphogenesis in the Early Zebrafish Embryo. PLoS Genet 2010, 6:e1000936.

42. Rizzoti K, Brunelli S, Carmignac D, Thomas PQ, Robinson IC, Lovell-Badge R: SOX3 is required during the formation of the hypothalamo-pituitary axis. Nat Genet 2004, 36:247–255.

43. Wegner M, Stolt CC: From stem cells to neurons and glia: a Soxist’s view of neural development. Trends Neurosci 2005, 28:583–588.

44. Collignon J, Sockanathan S, Hacker A, Cohen-Tannoudji M, Norris D, Rastan S, Stevanovic M, Goodfellow PN, Lovell-Badge R: A comparison of the properties of Sox-3 with Sry and two related genes, Sox-1 and Sox-2. Development 1996, 122:509–520.

45. McKimmie C, Woerfel G, Russell S: Conserved genomic organisation of Group B Sox genes in insects. BMC Genet 2005, 6:26.

46. Malas S, Duthie S, Deloukas P, Episkopou V: The isolation and high-resolution chromosomal mapping of human SOX14 and SOX21; two members of the SOX gene family related to SOX1, SOX2, and SOX3. Mamm Genome Off J Int Mamm Genome Soc 1999, 10:934–937.

47. Wegner M: SOX after SOX: SOXession regulates neurogenesis. Genes Dev 2011, 25:2423–2428.

48. Uchikawa M, Kamachi Y, Kondoh H: Two distinct subgroups of Group B Sox genes for transcriptional activators and repressors: their expression during embryonic organogenesis of the chicken. Mech Dev 1999, 84:103–120.

49. Uchikawa M, Yoshida M, Iwafuchi-Doi M, Matsuda K, Ishida Y, Takemoto T, Kondoh H: B1 and B2 Sox gene expression during neural plate development in chicken and mouse embryos: Universal versus species-dependent features. Dev Growth Differ 2011, 53:761–771.

50. Sanchez-Soriano N, Russell S: The Drosophila SOX-domain protein Dichaete is required for the development of the central nervous system midline. Development 1998, 125:3989–3996.

51. Ferrero E, Fischer B, Russell S: SoxNeuro orchestrates central nervous system specification and differentiation in Drosophila and is only partially redundant with Dichaete. Genome Biol 2014: R74.

52. Ambrosetti D-C, Basilico C, Dailey L: Synergistic activation of the fibroblast growth factor 4 enhancer by Sox2 and Oct-3 depends on protein-protein interactions facilitated by a specific spatial arrangement of factor binding sites. Mol Cell Biol 1997, 17:6321–6329.

53. Archer TC, Jin J, Casey ES: Interaction of Sox1, Sox2, Sox3 and Oct4 during primary neurogenesis. Dev Biol 2011, 350:429–440.

54. Bery A, Martynoga B, Guillemot F, Joly J-S, Retaux S: Characterization of Enhancers Active in the Mouse Embryonic Cerebral Cortex Suggests Sox/Pou cis-Regulatory Logics and Heterogeneity of Cortical Progenitors. Cereb Cortex 2013.

55. Cremazy F, Berta P, Girard F: Sox Neuro, a new Drosophila Sox gene expressed in the developing central nervous system. Mech Dev 2000, 93:215–219.

56. Ma Y, Certel K, Gao Y, Niemitz E, Mosher J, Mukherjee A, Mutsuddi M, Huseinovic N, Crews ST, Johnson WA, others: Functional Interactions between Drosophila bHLH/PAS, Sox, and POU Transcription Factors Regulate CNS Midline Expression of the slit Gene. J Neurosci 2000, 20:4596–4605.

57. Masui S, Nakatake Y, Toyooka Y, Shimosato D, Yagi R, Takahashi K, Okochi H, Okuda A, Matoba R, Sharov AA, Ko MSH, Niwa H: Pluripotency governed by Sox2 via regulation of Oct3/4 expression in mouse embryonic stem cells. Nat Cell Biol 2007, 9:625–635.

58. Tanaka S, Kamachi Y, Tanouchi A, Hamada H, Jing N, Kondoh H: Interplay of SOX and POU Factors in Regulation of the Nestin Gene in Neural Primordial Cells. Mol Cell Biol 2004, 24:8834–8846.

59. Wilson MJ, Dearden PK: Evolution of the insect Sox genes. BMC Evol Biol 2008, 8:120.

60. Zhong L, Wang D, Gan X, Yang T, He S: Parallel Expansions of Sox Transcription Factor Group B Predating the Diversifications of the Arthropods and Jawed Vertebrates. PLoS ONE 2011, 6:e16570.

61. Buescher M, Hing FS, Chia W: Formation of neuroblasts in the embryonic central nervous system of Drosophila melanogaster is controlled by SoxNeuro. Development 2002, 129:4193–4203.

62. Girard F, Joly W, Savare J, Bonneaud N, Ferraz C, Maschat F: Chromatin immunoprecipitation reveals a novel role for the Drosophila SoxNeuro transcription factor in axonal patterning. Dev Biol 299:530–542.

63. Sanchez-Soriano N, Russell S: Regulatory Mutations of the Drosophila Sox Gene Dichaete Reveal New Functions in Embryonic Brain and Hindgut Development. Dev Biol 2000, 220:307–321.

64. Shen SP, Aleksic J, Russell S: Identifying targets of the Sox domain protein Dichaete in the Drosophila CNS via targeted expression of dominant negative proteins. BMC Dev Biol 2013, 13:1.

65. Overton P: The role of Sox genes in the development of Drosophila melanogaster. University of Cambridge; 2003.

66. Overton PM, Meadows LA, Urban J, Russell S: Evidence for differential and redundant function of the Sox genes Dichaete and SoxN during CNS development in Drosophila. Development 2002, 129:4219–4228.

67. Aleksic J, Ferrero E, Fischer B, Shen SP, Russell S: The role of Dichaete in transcriptional regulation during Drosophila embryonic development. BMC Genomics 2013, 14:861.

68. Zhao G, Wheeler SR, Skeath JB: Genetic control of dorsoventral patterning and neuroblast specification in the Drosophila Central Nervous System. Int J Dev Biol 2007, 51:107–115.

69. Isshiki T, Pearson B, Holbrook S, Doe CQ: Drosophila neuroblasts sequentially express transcription factors which specify the temporal identity of their neuronal progeny. Cell 2001, 106:511.

70. Maurange C, Gould AP: Brainy but not too brainy: starting and stopping neuroblast divisions in Drosophila. Trends Neurosci 2005, 28:30–36.

71. Nowak MA, Boerlijst MC, Cooke J, Smith JM: Evolution of genetic redundancy. Nature 1997, 388:167–171.

72. Tautz D: Redundancies, development and the flow of information. BioEssays News Rev Mol Cell Dev Biol 1992, 14:263–266.

73. Wagner A: Gene duplications, robustness and evolutionary innovations. BioEssays 2008, 30:367–373.

74. Greer JM, Puetz J, Thomas KR, Capecchi MR: Maintenance of functional equivalence during paralogous Hox gene evolution. Nature 2000, 403:661–665.

75. Maconochie M, Nonchev S, Morrison A, Krumlauf R: Paralogous Hox genes: function and regulation. Annu Rev Genet 1996, 30:529–556.

76. Bradley RK, Li X-Y, Trapnell C, Davidson S, Pachter L, Chu HC, Tonkin LA, Biggin MD, Eisen MB: Binding Site Turnover Produces Pervasive Quantitative Changes in Transcription Factor Binding between Closely Related Drosophila Species. PLoS Biol 2010, 8:e1000343.

77. He Q, Bardet AF, Patton B, Purvis J, Johnston J, Paulson A, Gogol M, Stark A, Zeitlinger J: High conservation of transcription factor binding and evidence for combinatorial regulation across six Drosophila species. Nat Genet 2011, 43:414–420.

78. MacArthur S, Li X-Y, Li J, Brown JB, Chu HC, Zeng L, Grondona BP, Hechmer A, Simirenko L, Keranen SV: Developmental roles of 21 Drosophila transcription factors are determined by quantitative differences in binding to an overlapping set of thousands of genomic regions. Genome Biol 2009, 10:R80.

79. Paris M, Kaplan T, Li XY, Villalta JE, Lott SE, Eisen MB: Extensive Divergence of Transcription Factor Binding in Drosophila Embryos with Highly Conserved Gene Expression. PLoS Genet 2013, 9:e1003748.

80. Odom DT, Dowell RD, Jacobsen ES, Gordon W, Danford TW, MacIsaac KD, Rolfe PA, Conboy CM, Gifford DK, Fraenkel E: TissueVspecific transcriptional regulation has diverged significantly between human and mouse. Nat Genet 2007, 39:730–732.

81. Schmidt D, Wilson MD, Ballester B, Schwalie PC, Brown GD, Marshall A, Kutter C, Watt S, Martinez-Jimenez CP, Mackay S, Talianidis I, Flicek P, Odom DT: FiveVVertebrate ChIP-seq Reveals the Evolutionary Dynamics of Transcription Factor Binding. Science 2010, 328:1036–1040.

82. Stefflova K, Thybert D, Wilson MD, Streeter I, Aleksic J, Karagianni P, Brazma A, Adams DJ, Talianidis I, Marioni JC, Flicek P, Odom DT: Cooperativity and Rapid Evolution of Cobound Transcription Factors in Closely Related Mammals. Cell 2013, 154:530–540.

83. Arnold CD, Gerlach D, Spies D, Matts JA, Sytnikova YA, Pagani M, Lau NC, Stark A: Quantitative genomeVwide enhancer activity maps for five Drosophila species show functional enhancer conservation and turnover during cisVregulatory evolution. Nat Genet 2014.

84. Gallo SM, Gerrard DT, Miner D, Simich M, Des Soye B, Bergman CM, Halfon MS: REDfly v3.0: toward a comprehensive database of transcriptional regulatory elements in Drosophila. Nucleic Acids Res 2010, 39(Database):D118–D123.

85. Manning L, Heckscher ES, Purice MD, Roberts J, Bennett AL, Kroll JR, Pollard JL, Strader ME, Lupton JR, Dyukareva AV, Doan PN, Bauer DM, Wilbur AN, Tanner S, Kelly JJ, Lai S-L, Tran KD, Kohwi M, Laverty TR, Pearson JC, Crews ST, Rubin GM, Doe CQ: A Resource for Manipulating Gene Expression and Analyzing cis-Regulatory Modules in the Drosophila CNS. Cell Rep 2012, 2:1002¬1013.

86. Ross-Innes CS, Stark R, Teschendorff AE, Holmes KA, Ali HR, Dunning MJ, Brown GD, Gojis O, Ellis IO, Green AR, Ali S, Chin S-F, Palmieri C, Caldas C, Carroll JS: Differential oestrogen receptor binding is associated with clinical outcome in breast cancer. Nature 2012.

87. Ludwig MZ, Manu Kittler R, White KP, Kreitman M: Consequences of Eukaryotic Enhancer Architecture for Gene Expression Dynamics, Development, and Fitness. PLoS Genet 2011, 7:e1002364.

88. Perry MW, Boettiger AN, Bothma JP, Levine M: Shadow Enhancers Foster Robustness of Drosophila Gastrulation. Curr Biol 2010, 20:1562–1567.

89. Grant CE, Bailey TL, Noble WS: FIMO: scanning for occurrences of a given motif. Bioinformatics 2011, 27:1017–1018.

90. Quinlan AR, Hall IM: BEDTools: a flexible suite of utilities for comparing genomic features. Bioinformatics 2010, 26:841–842.

91. Löytynoja A, Goldman N: Phylogeny-Aware Gap Placement Prevents Errors in Sequence Alignment and Evolutionary Analysis. Science 2008, 320:1632–1635.

92. Löytynoja A, Goldman N: An algorithm for progressive multiple alignment of sequences with insertions. Proc Natl Acad Sci U S A 2005, 102:10557–10562.

93. Thomas-Chollier M, Defrance M, Medina-Rivera A, Sand O, Herrmann C, Thieffry D, van Helden J: RSAT 2011: regulatory sequence analysis tools. Nucleic Acids Res 2011, 39(suppl):W86–W91.

94. Turatsinze J-V, Thomas-Chollier M, Defrance M, van Helden J: Using RSAT to scan genome sequences for transcription factor binding sites and cis-regulatory modules. Nat Protoc 2008, 3:1578–1588.

95. Vogel MJ, Peric-Hupkes D, van Steensel B: Detection of *in vivo* protein-DNA interactions using DamID in mammalian cells. Nat Protoc 2007, 2:1467–1478.

96. Lee T, Marticke S, Sung C, Robinow S, Luo L: Cell-Autonomous Requirement of the USP/EcR-B Ecdysone Receptor for Mushroom Body Neuronal Remodeling in Drosophila. Neuron, 28:807–818.

97. Parrish JZ: Genome-wide analyses identify transcription factors required for proper morphogenesis of Drosophila sensory neuron dendrites. Genes Dev 2006, 20:820–835.

98. Kispert A, Herrmann BG, Leptin M, Reuter R: Homologs of the mouse Brachyury gene are involved in the specification of posterior terminal structures in Drosophila, Tribolium, and Locusta Genes Dev 1994, 8:2137–2150.

99. Murakami R, Takashima S, Hamaguchi T: Developmental genetics of the Drosophila gut: specification of primordia, subdivision and overt-differentiation. Cell Mol Biol Noisy–Gd Fr 1999, 45:661–676.

100. Hare EE, Peterson BK, Iyer VN, Meier R, Eisen MB: Sepsid even-skipped Enhancers Are Functionally Conserved in Drosophila Despite Lack of Sequence Conservation. PLoS Genet 2008: e1000106.

101. Clark AG, Eisen MB, Smith DR, Bergman CM, Oliver B, Markow TA, Kaufman TC, Kellis M, Gelbart W, Iyer VN, Pollard DA, Sackton TB, Larracuente AM, Singh ND, Abad JP, Abt DN, Adryan B, Aguade M, Akashi H, Anderson WW, Aquadro CF, Ardell DH, Arguello R, Artieri CG, Barbash DA, Barker D, Barsanti P, Batterham P, Batzoglou S, Begun D, et al.: Evolution of genes and genomes on the Drosophila phylogeny. Nature 2007, 450:203–218.

102. Stark A, Lin MF, Kheradpour P, Pedersen JS, Parts L, Carlson JW, Crosby MA, Rasmussen MD, Roy S, Deoras AN, Ruby JG, Brennecke J, curators HF, Project BDG, Hodges E, Hinrichs AS, Caspi A, Paten B, Park S-W, Han MV, Maeder ML, Polansky BJ, Robson BE, Aerts S, van Helden J, Hassan B, Gilbert DG, Eastman DA, Rice M, Weir M, et al.: Discovery of functional elements in 12 Drosophila genomes using evolutionary signatures. Nature 2007, 450:219–232.

103. Vavouri T, Semple JI, Lehner B: Widespread conservation of genetic redundancy during a billion years of eukaryotic evolution. Trends Genet, 24:485–488.

104. Wagner A: Distributed robustness versus redundancy as causes of mutational robustness. BioEssays 2005, 27:176–188.

105. Ferrari S, Harley VR, Pontiggia A, Goodfellow PN, Lovell-Badge R, Bianchi ME: SRY, like HMG1, recognizes sharp angles in DNA. EMBO J 1992, 11:4497.

106. Ghavi-Helm Y, Klein FA, Pakozdi T, Ciglar L, Noordermeer D, Huber W, Furlong EEM: Enhancer loops appear stable during development and are associated with paused polymerase. Nature 2014.

107. Russell SR, Sanchez-Soriano N, Wright CR, Ashburner M: The Dichaete gene of Drosophila melanogaster encodes a SOXVdomain protein required for embryonic segmentation. Development 1996, 122:3669–3676.

108. Overton PM, Chia W, Buescher M: The Drosophila HMG-domain proteins SoxNeuro and Dichaete direct trichome formation via the activation of shavenbaby and the restriction of Wingless pathway activity. Development 2007, 134:2807–2813.

109. Chan S-K, Ryoo H-D, Gould A, Krumlauf R, Mann RS: Switching the *in vivo* specificity of a minimal HoxVresponsive element. Development 1997, 124:2007–2014.

110. Slattery M, Ma L, Négre N, White KP, Mann RS: GenomeVWide TissueVSpecific Occupancy of the Hox Protein Ultrabithorax and Hox Cofactor Homothorax in Drosophila. PLoS ONE 2011, 6:e14686.

111. Coyne JA, Elwyn S, Kim SY, Llopart A: Genetic studies of two sister species in the Drosophila melanogaster subgroup, D. yakuba and D. santomea. Genet Res 2004, 84:11–26.

112. Riaz F: The application of mutant DNA adenine methyltransferase enzymes in Dam Identification. University of Cambridge; 2009.

113. Horn C, Wimmer EA: A versatile vector set for animal transgenesis. Dev Genes Evol 2000, 210:630–637.

114. Martin M: Cutadapt removes adapter sequences from highVthroughput sequencing reads. EMBnet J 2011, 17:pp-10.

115. Langmead B, Salzberg SL: Fast gappedVread alignment with Bowtie 2. Nat Methods 2012, 9:357–359.

116. Adams MD: The Genome Sequence of *Drosophila* melanogaster. Science 2000, 287:2185–2195.

117. Celniker SE, Wheeler DA, Kronmiller B, Carlson JW, Halpern A, Patel S, Adams M, Champe M, Dugan SP, Frise E: Finishing a wholeVgenome shotgun: release 3 of the *Drosophila* melanogaster euchromatic genome sequence. Genome Biol 2002, 3:research0079.

118. Richards S: Comparative genome sequencing of Drosophila pseudoobscura: Chromosomal, gene, and cisVelement evolution. Genome Res 2005, 15:1–18.

119. Li H, Handsaker B, Wysoker A, Fennell T, Ruan J, Homer N, Marth G, Abecasis G, Durbin R, 1000 Genome Project Data Processing Subgroup: The Sequence Alignment/Map format and SAMtools. Bioinformatics 2009, 25:2078–2079.

120. Heinz S, Benner C, Spann N, Bertolino E, Lin YC, Laslo P, Cheng JX, Murre C, Singh H, Glass CK: Simple Combinations of LineageVDetermining Transcription Factors Prime cisVRegulatory Elements Required for Macrophage and B Cell Identities. Mol Cell 2010, 38:576–589.

121. Love MI, Huber W, Anders S: Moderated estimation of fold change and dispersion for RNAVseq data with DESeq2. bioRxiv 2014.

122. Bardet AF, He Q, Zeitlinger J, Stark A: A computational pipeline for comparative ChIP-seq analyses. Nat Protoc 2011, 7:45–61.

123. Yu G ChlPseeker: ChlPseeker for ChIP Peak Annotation, Comparison, and Visualization. R; 2014.

124. Zhu LJ, Gazin C, Lawson ND, Pages H, Lin SM, Lapointe DS, Green MR: ChIPpeakAnno: a Bioconductor package to annotate ChIP-seq and ChIP-chip data. BMC Bioinformatics 2010, 11:237.

125. Nicol JW, Helt GA, Blanchard SG, Raja A, Loraine AE: The Integrated Genome Browser: free software for distribution and exploration of genomeVscale datasets. Bioinformatics 2009, 25:2730–2731.

126. Lyne R, Smith R, Rutherford K, Wakeling M, Varley A, Guillier F, Janssens H, Ji W, Mclaren P, North P: FlyMine: an integrated database for Drosophila and Anopheles genomics. Genome Biol 8:R129.

127. Edgar R, Domrachev M, Lash AE: Gene Expression Omnibus: NCBI gene expression and hybridization array data repository. Nucleic Acids Res 2002, 30:207–210.

